# Identification of B cell subsets based on antigen receptor sequences using deep learning

**DOI:** 10.1101/2024.02.06.579098

**Authors:** Hyunho Lee, Kyoungseob Shin, Yongju Lee, Soobin Lee, Seungyoun Lee, Eunjae Lee, Seung Woo Kim, Ha Young Shin, Jong Hoon Kim, Junho Chung, Sunghoon Kwon

## Abstract

B cell receptors (BCRs) denote antigen specificity, while corresponding cell subsets indicate B cell functionality. Since each B cell uniquely encodes this combination, physical isolation and subsequent processing of individual B cells become indispensable to identify both attributes. However, this approach accompanies high costs and inevitable information loss, hindering high-throughput investigation of B cell populations. Here, we present BCR-SORT, a deep learning model that predicts cell subsets from their corresponding BCR sequences by leveraging B cell activation and maturation signatures encoded within BCR sequences. Subsequently, BCR-SORT is demonstrated to improve reconstruction of BCR phylogenetic trees, and reproduce results consistent with those verified using physical isolation-based methods or prior knowledge. Notably, when applied to BCR sequences from COVID-19 vaccine recipients, it revealed inter-individual heterogeneity of evolutionary trajectories towards Omicron-binding memory B cells. Overall, BCR-SORT offers great potential to improve our understanding of B cell responses.

## Introduction

From infectious diseases to various types of cancers and autoimmune diseases, B cells play a crucial role in establishing antibody-based protective immunity(1–7). Within the highly heterogeneous B cell populations, individual B cells play distinct roles, each expressing a unique combination of cell subset and antigen receptor(8–10). The cell subset serves as a primary criterion for the functionality of B cells, on the other hand, the BCR is an antibody molecule uniquely expressed by each B cell to recognize its cognate antigen. Given that the BCR and the corresponding cell subset represent two different modalities of the B cell, simultaneous interrogation of both is essential for comprehensive understanding about B cells.

Before antigenic exposure, the BCRs expressed by naïve B cells establish the foundation for preexisting immunity, preparing the initiation of the B cell response(11,12). When faced with an immunological challenge, B cells undergo a maturation process to optimize immunological role of the B cells by developing their cell subsets and BCR sequences simultaneously. In detail, naïve B cells differentiate into either memory B cells or antibody-secreting cells (ASCs), while at the same time, the BCR sequences also mature through somatic hypermutations (SHMs) and class switching(13). Notably, this bifurcation of maturation trajectories carry distinct implications. BCRs expressed by memory B cells epitomize past antigenic encounters, thus widely utilized in identifying immunological memories engraved in vaccine recipients or convalescent patients. For example, the human antibody response against SARS-CoV-2 was revealed by the discovery of memory B cells expressing neutralizing BCRs among convalescent COVID-19 patients(14–19). On the other hand, BCRs expressed by ASCs reflect the current antigenic challenges, thus widely utilized in circumstances under explicit exposure to antigens, such as autoimmune disorders or ongoing severe infections. As an instance, potential autoreactive BCRs and their maturations through continuous exposure to autoantigens were analyzed by examining ASCs persistent in autoimmune disease patients(20). Collectively, coupling of antigen receptor and its originating cell subset resolves the role of B cells while unveiling the maturation trajectories.

Various methods have been utilized to investigate B cell subset and antigen receptor. However, approaches combining B cell subset and antigen receptor information of individual B cells while addressing the high diversity of B cell populations are lacking. For example, fluorescence-activated cell sorting (FACS)(21–25) or single B cell RNA sequencing (scRNA-seq)(4–7,9) was utilized to identify B cell subset and antigen receptor simultaneously. However, both methods require additional complex machinery and experimental procedures for the physical isolation and processing of numerous bulk B cells in a single-cell resolution, which is costly and susceptible to information loss(26). In detail, compartmentalization of multiple cell subsets using FACS necessitates consecutive gating rounds and the preparation of separate libraries for sequencing. Therefore, in most cases, practical use is constrained to specific cell subsets of interest while excluding the other cell subsets(27). This results in a loss of information regarding the transition landscape between cell subsets, which is inherent in most B cell maturation processes. On the other hand, scRNA-seq couldn’t cover the vast diversity of the entire B cell population owing to prohibitive expenses, thus resulting in severe undersampling of information. In practice, scRNA-seq covers B cell diversity that is more than one order of magnitude lower compared to bulk BCR high-throughput sequencing (HTS), thereby erasing the maturational relationships between B cells.

Prediction of cell subsets directly from the sequence of corresponding antigen receptors could provide high throughput information of antigen receptors combined with their cell types. In previous studies, SHMs induced in the IGHV region were quantified and B cells exhibiting low SHMs and un-switched isotypes (IgM/IgD) were categorized as naïve B cells, while the rest were classified as antigen-experienced B cells (either memory B cells or ASCs)(28–30). Yet, this heuristic approach was unable to distinguish between memory B cells and ASCs. Considering that the complementarity-determining region 3 of heavy chain (HCDR3) region undergoes the most frequent SHMs, and that B cell activation and maturation are driven by the binding of its BCR to a cognate antigen, there is a chance that HCDR3 harbors pivotal information for predicting cell types. However, no method has yet been proposed to exploit the potential of HCDR3 information. Recently, a machine learning approach was proposed, which harnessed HCDR3 information for cell subset prediction(23). However, fixed vector embeddings that represent physicochemical properties of amino acids were inadequate for HCDR3 sequence representations, resulting in a performance suboptimal to replace FACS or scRNA-seq. Thus, a novel method to utilize HCDR3 as a valuable source of information for cell subset prediction is essential.

To this end, we proposed BCR-SORT, a deep learning model that predicts the originating B cell subset using the HCDR3 sequence of a given BCR. Unlike previous approaches, BCR-SORT established a direct link between the BCR and its originating cell subset by deciphering the inherent B cell maturation features encoded within the HCDR3 sequence. Exploiting these features, BCR-SORT offered a scalable and cost-effective method to accurately couple antigen receptors with their cell subsets, especially when used in conjunction with HTS of the BCR repertoire (Figure 1A). Through benchmark tests against FACS and scRNA-seq, we validated the applicability of BCR-SORT on datasets obtained from diverse immunological conditions. In addition, we demonstrated that BCR-SORT enabled cell subset-aware reconstruction of BCR lineage, which rearranged the original evolutionary scenario of the lineage to follow the biological process of B cell differentiation (Figure 1B). Finally, BCR-SORT was applied to various unlabeled datasets from autoimmune diseases and vaccinations. Of note, BCR-SORT revealed treatment-resistant ASC populations from autoimmune disease patients, and at the same time, revealed the maturation trajectory towards Omicron-binding memory B cells induced by the triple vaccinations of wild-type SARS-CoV-2 virus.

**Figure 1.**
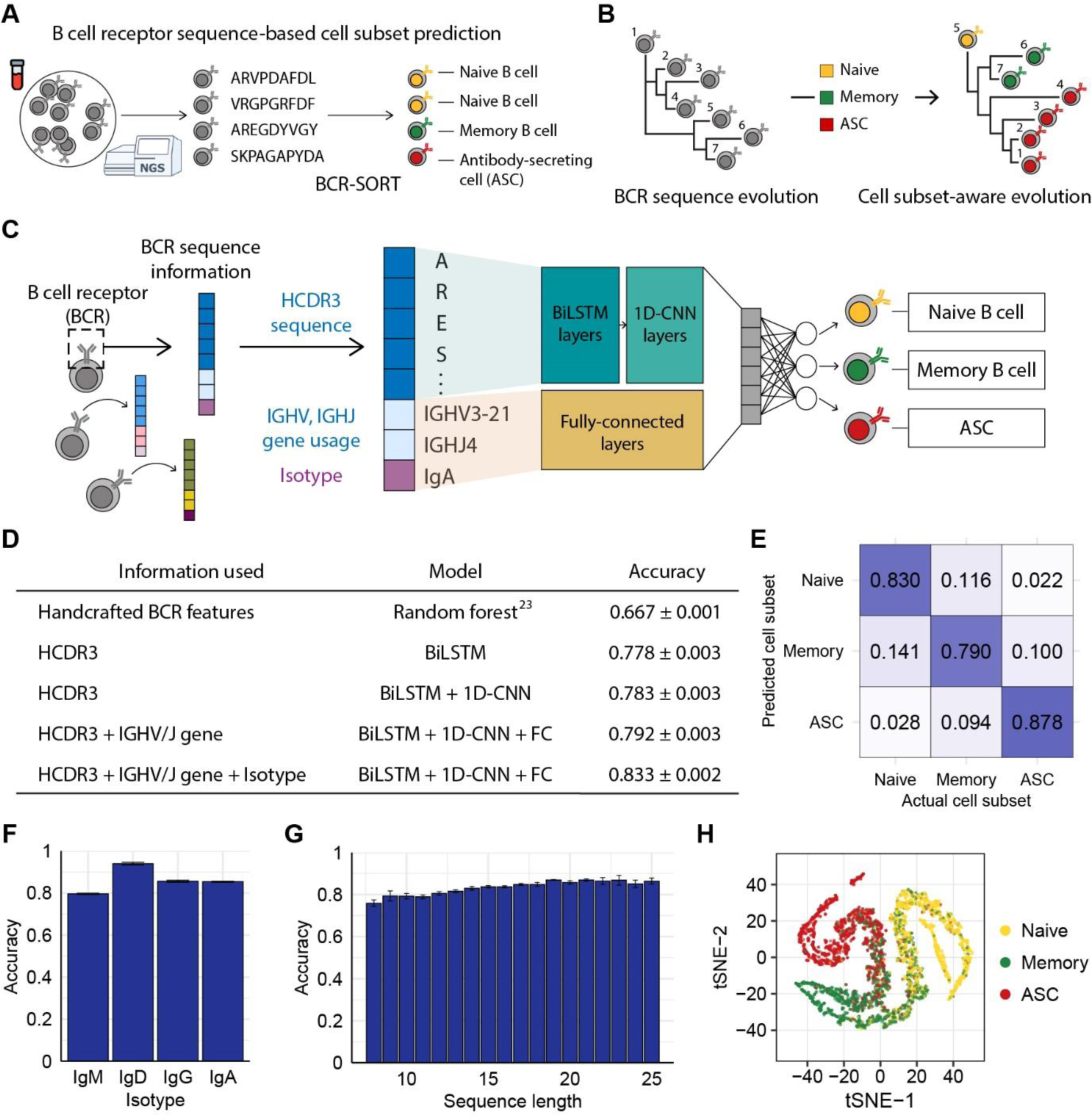
BCR sequence-based prediction of B cell subsets using BCR-SORT. (**A**) Schematic depicting the coupling of B cell subsets with corresponding antigen receptors using BCR-SORT. (**B**) Schematic example of BCR lineage rearrangement using BCR-SORT. Lineage inferred without cell subset information is reconstructed using BCR-SORT. (**C**) Detailed illustration of the BCR-SORT architecture. (**D**) Performance of BCR-SORT with respect to input attributes. The influence of input attributes on accuracy is shown, including that in the case of the existing state-of-the-art method. (**E**) Confusion matrix representing the average prediction results of BCR-SORT with respect to B cell subsets. (**F, G**) Bar plots representing the relationship between model accuracy and the isotype of input instances (**F**), and HCDR3 sequence length (**G**), respectively. (**H**) tSNE visualization of penultimate layer activation in BCR-SORT. 5,000 instances are randomly selected. Accuracy in (**D**-**G**) is measured with 5-fold cross validation.

## Materials and Methods

### Model architecture and training

BCR-SORT was constructed to process HCDR3 sequence, IGHV gene, IGHJ gene, and isotype as inputs and conducted multi-class classification. HCDR3 amino acid sequences were converted to numeric feature vectors by first encoding them using learnable embedding vectors and then processing them using 2-layer bidirectional Long Short-Term Memory (LSTM) followed by 3-layers of 1-dimensional Convolutional Neural Network (1D-CNN). Other input attributes were also converted to embedding vectors and concatenated with the sequence feature vector. The concatenated feature vectors were fed into a 3-layer multi-layer perceptron (MLP) to produce output vector. For better generalizability, we randomly masked a single amino acid from each HCDR3 sequence and created an auxiliary task to predict the masked amino acid using the concatenated feature vector. Total loss was calculated by the sum of cross entropy loss for the original classification task and the auxiliary task after scaling the auxiliary loss to 0.05. The model was trained using a learning rate of 10^-4^ and a batch size of 1024. The accuracy of the model was assessed by 5-fold cross validation. PyTorch (v1.12.0) was used for the implementation of LSTM, CNN, and MLP.

### Dataset preparation

We used the BCR-B cell subset coupled data archived in the Observed Antibody Space(31) (OAS) database to train BCR-SORT. We confirmed that each dataset was constructed while satisfying the following criteria: naïve B cell (CD19+CD27-), memory B cell (CD19+CD27+CD38-), and ASC (CD19+CD27+CD38+). Memory B cell data were further obtained from Mitsunaga et al.(25) for the sake of balancing with other labels. To identify HCDR3 sequence, IGHV, IGHJ, and isotype from each BCR sequence, sequence annotation results provided by OAS were utilized as input, whereas the data obtained from Mitsunaga et al. were annotated in-house using BLAST(32) and IgBLAST(33). HCDR3 sequences with length between 8 and 25 were used. For each B cell subset, the isotype proportion was balanced to reflect the actual proportion in the body, following a previous study(24) (Supplementary Table 4). Identical BCR sequences duplicated in multiple B cell subsets were discarded. In total, 2.65 million sequences was collected and split into training set (98%), validation set (1%) and test set (1%).

To evaluate the performance of the state-of-the-art model proposed in previous work(23), additional BCR sequence features (the number of mutations, the physicochemical properties of junction sequences, and the replacement/silent mutation ratios) were calculated in R using shazam(34) (v1.1.2) and alakazam(34) package (v1.2.1).

### Integrated Gradients (IG)

IG is a model interpretation method that calculates the cumulative sum of gradients along a path from a baseline value to the input value to measure the influence score of each input feature. Each input consisted of 25 amino acid sequence features (including padding), 1 isotype feature, 1 IGHV gene usage feature, and 1 IGHJ gene usage feature, and the IG value was calculated with regard to the baseline feature. We used the zero vector as a baseline feature, which was also an embedding of the padding token. The IG value was calculated as follows.

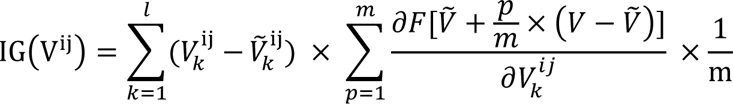

where *F* denotes the BCR-SORT model, *V* denotes an input feature, 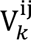 denotes *k*-th value of the i-th input and j-th feature, *Ṽ* denotes a baseline feature, *p* denotes an interpolation step, and *m* denotes the number of total interpolation steps. As the IG value was calculated for each vector element, the final IG value for each input feature was computed as the average of the IG vector elements. To focus on features with high influence on the output, input features with top 10% IG values (high-IG) were selected for further analysis. IntegratedGradients function in the captum(35) (v0.5.0) module was used to calculate the IG values.

### Parapred for paratope prediction

Parapred is a sequence-based paratope prediction model that utilizes a bidirectional LSTM and CNN architecture to leverage local and neighboring residue features(36). By applying the pre-trained Parapred model to HCDR3 sequences, the probabilities of each residue corresponding to a paratope were obtained. Each amino acid residue was predicted as a paratope if the obtained probability was higher than 0.488, which was the threshold used in the original paper(36).

### *In silico* saturation mutagenesis

Using high-IG HCDR3 features, we applied *in silico* saturation mutagenesis by replacing the amino acid of each high-IG residue with 19 different amino acids, while retaining the other input features (isotype, IGHV gene usage, and IGHJ gene usage). To mimic the mechanism of SHM, sequence length was identically controlled and a single substitution of amino acid was mutated, since indels are rare in SHM(37). Subsequently, we predicted cell subsets of the mutated sequences using BCR-SORT and verified whether the applied mutation altered the prediction.

### Validation of *in silico* cell subset alteration on human repertoire

*In silico* mutations driving the alteration of B cell subset prediction was validated in *in vivo* circumstances using a human repertoire dataset. To reflect the actual *in vivo* alteration of B cell subsets, only human repertoires comprising the entire cell subsets (naive, memory, and ASC) in the identical dataset were used. We verified whether the original sequence and the mutated sequence pairs analyzed in *in silico* experiments were identifiable in the human repertoire as identical cell subsets. Each mutation was grouped based on the types of previous amino acids and the location of the mutation. Subsequently, the number of cell subset alterations for each mutation group was quantified based on the number of BCR sequences that induced a cell subset alteration via a single mutation within that mutation group. Finally, the correlation between mutation groups observed in *in vivo* alterations and *in silico* alterations was analyzed.

### Benchmarking FACS and scRNA-seq

External validation datasets for benchmarking FACS and scRNA-seq were prepared by manually downloading, and BCR sequences were annotated following the same manner with training set (Supplementary Table 3). In case of benchmarking FACS, BCR sequencing data acquired after sorting the single biological sample into three different B cell subsets using FACS was collected to evaluate the model in real-world settings. B cell subset sorted by the phenotypic criteria identical to the training set was utilized. In case of benchmarking scRNA-seq, each paper independently identified B cell subsets from their gene expression data, while we followed the annotation of B cell subset provided in each paper. In case of benchmarking FACS, the validation datasets were constructed per individual, while those from multiple individuals were pooled when benchmarking scRNA-seq due to data scarcity issue.

### Transfer learning

Trained BCR-SORT was fine-tuned into a disease-specific model or a chronological model using external datasets, which were prepared for benchmarking FACS and scRNA-seq. For fine-tuning, the model was initialized with the pre-trained weights and then further trained on the same task to achieve finely adjusted model. Corresponding to each type of disease, data from a single individual were provided for fine-tuning, and the accuracy on datasets from other individuals was measured to assess the model’s performance in unseen situations with data-scarcity. As data from multiple individuals were pooled in case of scRNA-seq, data from a single paper were used for fine-tuning and the accuracy on datasets from other papers with same diseases was measured. For chronological transfer, datasets containing multiple time points from identical individuals were selected. Similarly, data from a single time point were provided for fine-tuning, and the accuracy on datasets from different time points was measured for each individuals. All parameters were updated during fine-tuning without freezing specific layer to optimize the model(38), and auxiliary task was not provided during fine-tuning to prevent over-fitting on limited dataset. A learning rate of 0.1 times lower than the original learning rate was used to delicately adjust the model’s weights, ensuring that the pre-existing knowledge was not abruptly overridden by new data.

### Reconstruction of BCR lineage using phylogenetic analysis

Full-length nucleotide sequences in the BCR repertoire were processed for phylogenetic analysis following the workflow of Change-O(34). Briefly, BCR sequences sharing identical IGHV/IGHJ genes were assigned to a lineage when the similarity between junction sequences exceeded 85%. For each lineage, the virtual germline sequence was inferred after masking the IGHD segment due to its unreliable alignment. The phylogenetic tree of the BCR lineage was constructed using the BuildTrees method in IgPhyML(39,40). To consider lineages undergone active maturations, those containing 10 or more BCR sequences and isotype switching were taken into consideration. Lineages containing more than 200 BCR sequences were down-sampled to prevent excessive computation time required when constructing the phylogenetic tree(41). The phylogenetic tree was analyzed and visualized in R using dowser(42,43) (v1.1.0), and ggtree(44) (v3.6.2) package.

### Rerooting of BCR phylogenetic tree using BCR-SORT

Given a phylogenetic tree, we predicted cell subsets of constituent BCRs using BCR-SORT and examined whether antigen-experienced B cell (memory B cell or ASC) preceded antigen-unexperienced naïve B cell along the tree. If this was the case, we selected the least mutated naïve B cell, considering mutations induced within the entire IGHV region, as an alternative root of the tree, and rearranged the tree (“reroot”) based on the new root. IgPhyML was utilized to construct the rerooted phylogenetic tree, after removing the existing virtual germline and setting the new root of the tree as decided. In case of longitudinal studies providing lineages comprised of BCRs across multiple time points, the root was determined among BCRs obtained from the earliest time point in the lineage to reflect chronological development simultaneously.

### Blood sample collection

Peripheral blood samples were obtained at three time points from one patient with pemphigus vulgaris (PV) – at diagnosis, one month after the second dose of rituximab (RTX), and at relapse – and from one patient with myasthenia gravis (MG) – before RTX, 1 week after the first dose of RTX, and three months after RTX. The study involving human participant was approved by the Institutional Review Board of Gangnam Severance Hospital (IRB No. 3-2019-0191, PV) and by the Institutional Review Board of Severance Hospital, Yonsei University Health System (IRB No. 4-2023-1059, MG). Peripheral blood mononuclear cells (PBMCs) were obtained from blood samples of the patients using Ficoll–Paque (GE Healthcare). Total RNA was isolated using TRIzol Reagent (Invitrogen) according to the manufacturer’s protocols.

### Library preparation and next-generation sequencing (NGS)

Genes encoding the variable domain (VH) and part of the first constant domain of the heavy chain (CH1) were amplified, using specific primers, as described previously(11). Briefly, total RNA was used as a template to synthesize complementary DNA (cDNA), using the SuperScript IV First-Strand Synthesis System (Invitrogen), with specific primers targeting the CH1 domain of each isotype (IgM, IgD, IgG, IgA, and IgE), according to the manufacturer’s instructions. After cDNA synthesis, SPRI beads (Beckman Coulter, AMPure XP) were used to purify cDNA. The purified cDNA was subjected to second-strand synthesis using V gene-specific primers and KAPA Biosystems kit (Roche, KAPA HiFi HotStart PCR Kit). The PCR conditions for sample collected from the PV patient at one month after the second dose of RTX was as follows: 95°C for 3min; 4 cycles of 98°C for 30s, 60°C for 45s, 72°C for 1 min; and 72°C for 5 min. The PCR conditions for other samples were as follows: 95°C for 3min, 1 cycle of 98°C for 30s, 60°C for 45s, 72°C for 1 min, and 72°C for 5 min. After second-strand synthesis, double-stranded DNA was purified using SPRI beads. VH-CH1 genes were amplified using purified dsDNA using primers containing indexing sequences and a KAPA Biosystems kit. The PCR conditions were as follows: 95°C for 5 min; 25 cycles of 98°C for 30 s, 60°C for 30 s, and 72°C for 1 min; and 72°C for 5 min. PCR products were subjected to electrophoresis on a 1.5% agarose gel and purified using a QIAquick gel extraction kit (QIAGEN) according to the manufacturer’s instructions. The gel-purified PCR products were purified again using SPRI beads. SPRI-purified libraries were quantified with a 4200 TapeStation System (Agilent Technologies) using a D1000 ScreenTape assay and subjected to next-generation sequencing on the Illumina NovaSeq platform.

### NGS data processing

NGS data were processed following the procedures described previously(11). Briefly, NGS reads were merged by PEAR(45) (v0.9.10), and those with 95% of the bases having Phred scores higher than 20 were left for use. VH-CH1 primers and unique molecular identifier (UMI) sequences were recognized from each read, and the reads were clustered according to the UMI sequences. The clustered reads were aligned using Clustal Omega(46,47) (v1.2.4), and a consensus sequence was extracted within each UMI cluster based on the most dominant base pairs at each position. Isotype and V(D)J regions were annotated in-house for each consensus read, as described above.

### Persistent clone

Within the BCR repertoire from the PV patient, a BCR clone was defined as a group of BCR sequences sharing identical IGHV/IGHJ gene and HCDR3 amino acid sequence, following the definition of previous work(20). Clones present from pre-RTX to relapse were defined as persistent clones. Those comprised solely of IgM or IgD BCRs were excluded to focus on B cells eliciting antigen-specific responses.

### Identification of SARS-CoV-2-specific BCR lineage

BCR repertoire data provided by Park et al.(48) were processed to reconstruct the lineage, as described above. SARS-CoV-2-specific BCR sequences in the repertoires were identified using verified binder sequences archived in CoV-AbDab(49). Omicron-specific BCR sequences and the effective concentration (EC50) of BCRs within the Omicron binder lineages were identified from the data provided in the original paper. BCR sequences in the repertoire were labeled as binders based on the HCDR3 sequence by allowing a single amino acid mismatch.

To examine the relationship between consecutive vaccinations and Omicron immunity, lineages containing the verified Omicron binders after the 3^rd^ dose of vaccine were selected. To focus on vaccine antigen-driven responses, one naïve B cell lineage was discarded and eight different lineages were identified as a result. Among those eight lineages, memory B cell lineages (n=5) were subjected to phylogenetic analysis, considering the significance of memory B cells on driving Omicron immunity.

## Results

### Prediction of B cell subsets using BCR-SORT

We assumed that the BCR sequence, especially the HCDR3 sequence, possesses learnable features since activation and maturation of B cells are driven by the binding of their BCR with cognate antigens. Utilizing inputs comprised of HCDR3 amino acid sequence, IGHV/IGHJ gene usage, and isotype information, BCR-SORT predicted the most probable originating B cell subset among naïve B cell, memory B cell, and ASC, as those three cell subsets comprise the majority of the B cell population(50) (Figure 1C). BCR-SORT was trained on 2.6 million BCR sequences obtained from individuals with heterogeneous immunological conditions (Supplementary Table 1). By leveraging the HCDR3 sequence features extracted from LSTM layers followed by 1D-CNN layers, BCR-SORT predicted corresponding cell subsets with high accuracy, outperforming the existing state-of-the-art method based on handcrafted features(23) (Figure 1D). Although those handcrafted features were created using the entire variable regions of BCRs, BCR-SORT exhibited superior performance using only the HCDR3 sequences, indicating that HCDR3 inherently contains plenty of information to represent cell subsets.

In addition to the HCDR3 sequence, other input attributes also contributed to accurate prediction, with the isotype and IGHV/IGHJ gene following the HCDR3 sequence in terms of importance (Supplementary Table 2). Further, the performance of BCR-SORT was assessed over the entire set of cell subsets, isotypes, and HCDR3 lengths (Figures 1E-G). Interestingly, an increase in input sequence length was observed to enhance accuracy, further corroborating the benefits of HCDR3 information for prediction. Meanwhile, BCR sequences embedded by the BCR-SORT exhibited trajectory-like patterns spanning the entire B cell subsets, resembling the biological course of B cell differentiation as well as the actual cell sorting process (Figure 1H).

### Interpretation of BCR-SORT

Learning the relationship between BCRs and their corresponding cell subsets implies understanding the BCR sequence features relevant to B cell activation and differentiation. To ascertain whether BCR-SORT generated outputs by truly utilizing these features, we probed the relationships between input features and model predictions using Integrated Gradients (IG)(51). Through this method, we calculated the influence of each input feature on model output, and to focus on input features with high influence on the output, those with top 10% IG values (high-IG) were selected for further analysis.

Depending on B cell subsets, input features contributed to the prediction results to different extents (Figure 2A). Isotype features exhibited substantial influence when predicting naïve B cells as their isotypes are dominated by IgM and IgD in nature(25). Nevertheless, high-IG features corresponded predominantly to the HCDR3 sequence throughout the entire B cell subsets. Notably, the influence of HCDR3 was more pronounced in antigen-experienced cells (memory B cells and ASCs) than antigen-unexperienced naïve B cells, as BCR-SORT was found to be capable of utilizing B cell activation and maturation signatures encoded in the HCDR3 sequence.

**Figure 2.**
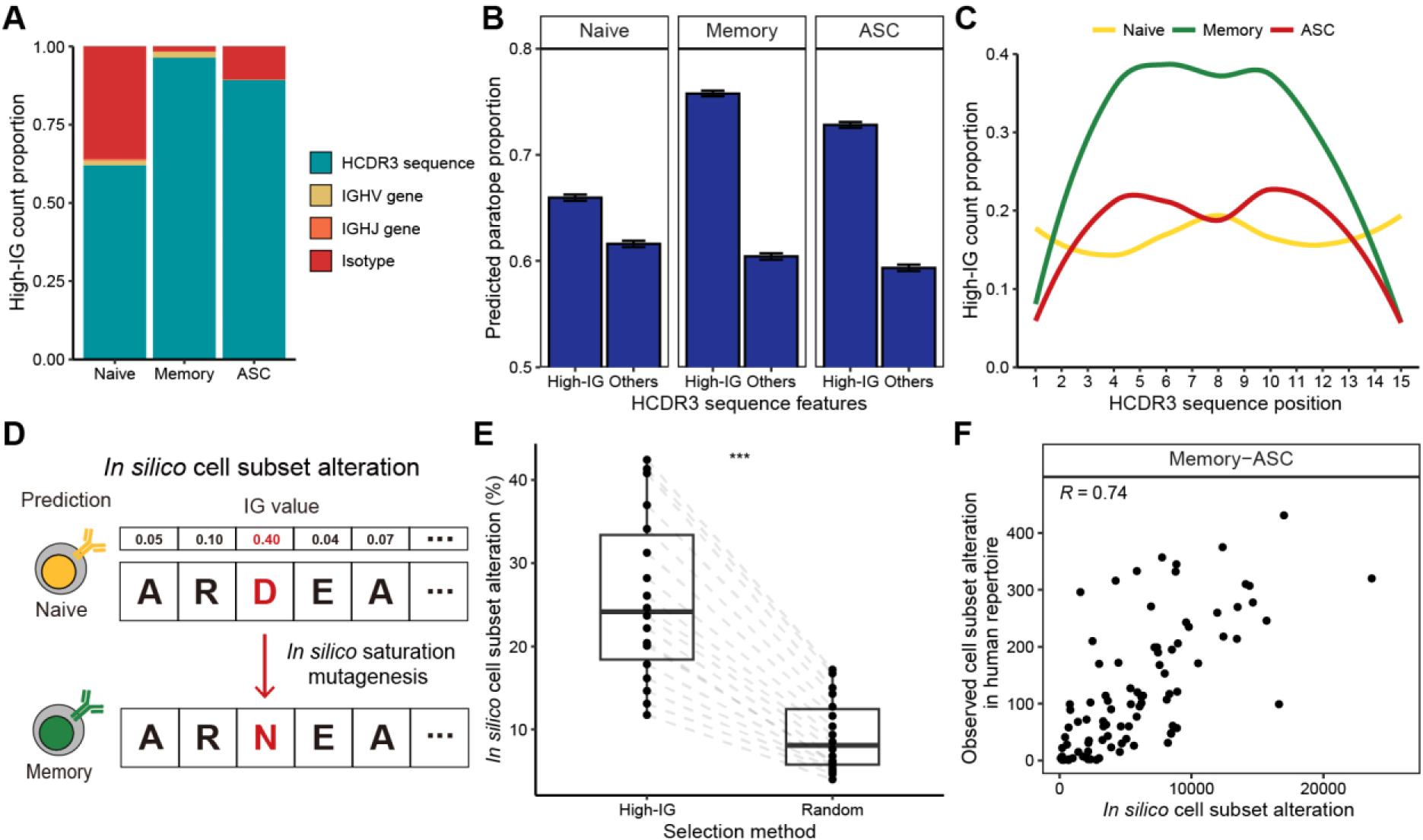
Interpretation of B cell subset-specific BCR sequence features using Integrated Gradients. (**A**) Composition of the top 10% high-IG features with regard to cell subsets. (**B**) Proportion of HCDR3 residues predicted as paratope using Parapred. Mean predicted paratope proportions and their confidence intervals (error bars) are plotted. The confidence interval is obtained by bootstrapping 100,000 high-IG and non-high-IG sequences 10,000 times. (**C**) Proportion of high-IG HCDR3 residues with regard to position. Sequence length of 15, which is the most dominant length in the dataset, is shown as an example. (**D**) Schematic of the *in-silico* saturation mutagenesis experiment. A single substitution of HCDR3 residue is considered in the region-of-interest, and B cell subset of the mutated sequence is predicted by BCR-SORT. (**E**) Proportion of *in silico* mutations altering the prediction of BCR-SORT. To examine if the mutation in high-IG residue was critical on altering the prediction, HCDR3 residues to induce mutations were selected from high-IG residues and randomly selected residues. The proportion is calculated with respect to sequence length. (**F**) Correlation plot between the cell subset alteration counts measured in *in silico* saturation mutagenesis and in human repertoire data. Alteration between memory B cell and ASC with the sequence length of 15 is plotted as an example. Spearman’s correlation coefficient is shown in the figure. In (**E**), the statistics are calculated using paired t-test. \*\*\**p* < 0.001.

B cell activation and maturation are associated with BCR-antigen binding and the accumulation of SHMs. In accordance with this, we discovered that high-IG values were assigned to HCDR3 residues related to BCR-antigen binding and SHM events. By identifying HCDR3 residues contributing to antigen binding (paratope) using Parapred(36), we observed that the proportion of paratope was significantly higher for high-IG residues (Figure 2B). This tendency was notable in antigen-experienced cells, indicating that BCR-SORT considered the antigen binding capability of BCRs during prediction. In addition, high-IG values were assigned to the middle part of the HCDR3, which encoded higher diversity owing to frequent SHMs compared to the amino-terminal and the carboxyl-terminal parts of the HCDR3(52) (torso of HCDR3), during the prediction of antigen-experienced cells (Figures 2C and S1). In sum, the aforementioned evidences indicated that the activation and maturation signatures encoded in BCR sequences were exploited by BCR-SORT.

Finally, we assessed the impact of high-IG residues on cell subset prediction. This was achieved by conducting *in silico* saturation mutagenesis on high-IG residues and identifying alterations in cell subset prediction(53). This experiment efficiently covered all possible mutation scenarios, while mimicking the mechanism of SHM (Figure 2D). Consequently, *in silico* mutations of a single high-IG residue were found to alter the model’s predictions with significantly higher probability than those induced in random residues (Figure 2E). While our primary focus was on *in silico* experiments, we found reproducible results in human BCR repertoire data. This was evident as identical mutations frequently observed in the *in silico* experiment were also repeatedly identified in the human BCR repertoire data (Figures 2F, S2).

### Benchmarking FACS and scRNA-seq

BCR-SORT was further validated by comparing the cell subset prediction results with cell subsets identified by FACS and scRNA-seq using external datasets. Assessed on multiple unseen datasets comprising various diseases to benchmark both FACS and scRNA-seq(4–7,21– 23,54) (Supplementary Table 3), BCR-SORT outperformed the current state-of-the-art method (Figures 3A, B). Although the BCR-SORT was trained on datasets originated from various diseases (Supplementary Table 1), it had no experience of datasets such as COVID-19, tetanus toxoid (TT) vaccination, and systemic lupus erythematosus (SLE). However, these unseen diseases were also included to assess the model’s general applicability in diverse physiological and pathological circumstances. Overall, BCR-SORT exhibited stable performance on various diseases, except a slight performance degradation on the SLE dataset.

**Figure 3.**
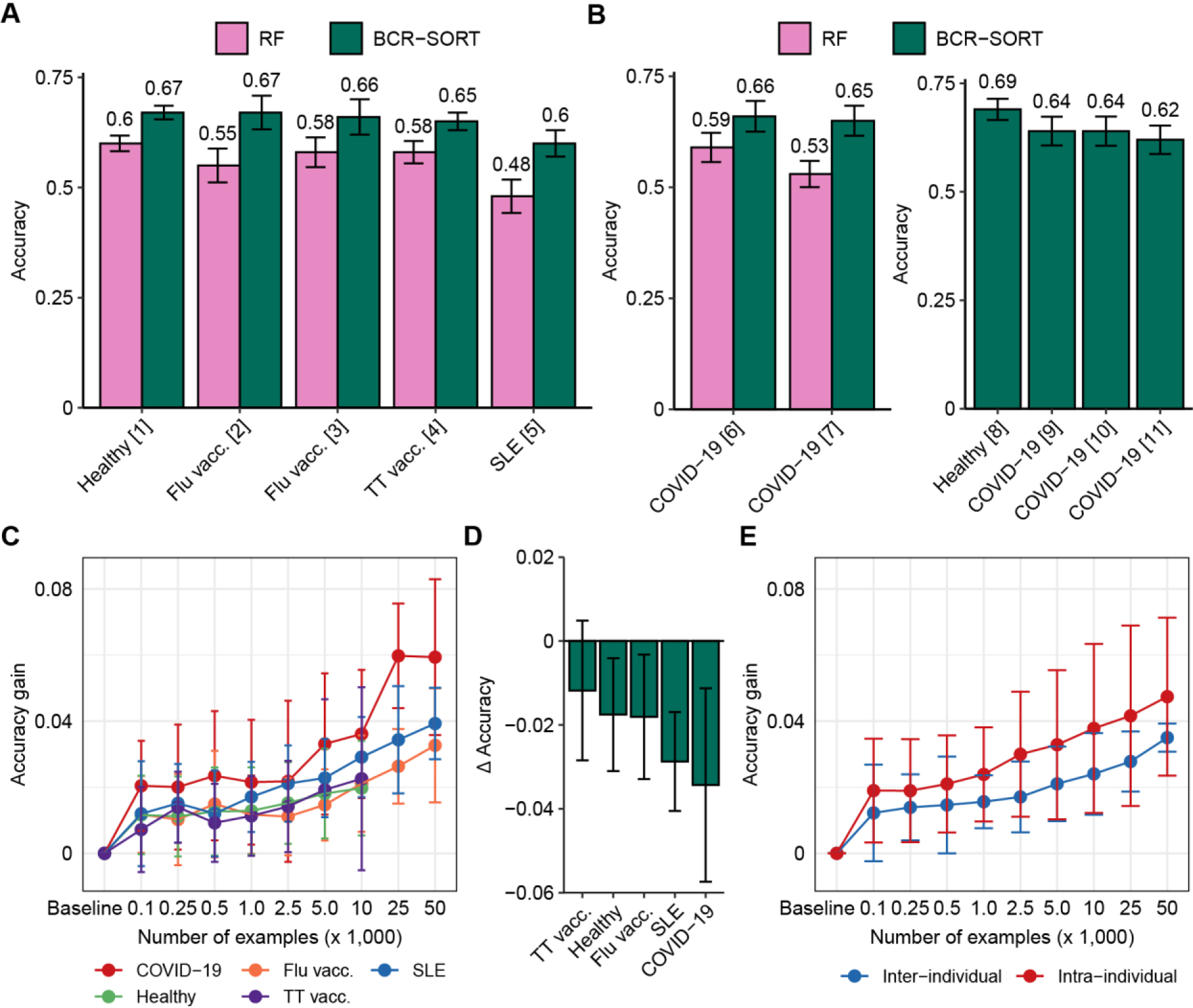
External validation and transfer learning for benchmarking FACS and scRNA-seq. (**A**, **B**) Performance of BCR-SORT and the current state-of-the-art random forest (RF)-based model on external validation datasets, verified by FACS (**A**) and scRNA-seq (**B**). Accuracy is measured using various benchmark datasets and presented according to the source of the dataset. Performance of RF is evaluated on datasets for whom full length of nucleotide BCR sequences are available, and error bars of datasets containing only a single sample are calculated by bootstrapping 150 BCR sequences 1,000 times. (**C**) Accuracy gains achieved by fine-tuning the established BCR-SORT into disease-specific models with respect to the number of examples utilized in fine-tuning. (**D**) Performance gap between models trained with the same disease-specific examples, but built either from the established BCR-SORT or from scratch. In cases of using maximum number of training examples are shown. (**E**) Comparison of accuracy gains achieved using two different data transfer schemes. In case of inter-individual transfer, the established BCR-SORT was fine-tuned using data from other individuals under identical type of disease (in the same manner with (**C**)), while in case of intra-individual transfer, data from identical individual’s chronological data was given.

In fact, BCR profiles expressed by each B cell subset are divergent depending on the type of disease owing to different antigen-specific B cell responses, which affects the prediction performance. We hypothesized that the performance of BCR-SORT can be further improved if it is optimized for specific diseases. Based on this hypothesis, we utilized transfer learning to overcome the differences originating from various diseases and leverage the full capacity of the model. Thus, for each type of disease, we fine-tuned the established BCR-SORT into disease-specific models and measured the improvements in accuracy (Figure 3C). Clearly, fine-tuning improved the model performance across all diseases by integrating specific disease-related features with the general cell subset features. By contrast, BCR-SORT trained from scratch using only the disease-specific data could not outperform these fine-tuned models (Figure 3D). Given that the type of disease is specified in most analyses and considering the effectiveness of disease-specific fine-tuning even in data-scarce settings, this approach can be viewed as generally applicable in practice.

In addition to disease-specific fine-tuning, BCR-SORT could be fine-tuned on single time point data of an individual to mitigate individual BCR heterogeneity. As the type of disease is likely to be consistent in most patients over multiple time points, we expected that the addition of individual-level BCR properties over disease-specific properties by chronological transfer would provide further accuracy. Despite the chronological gap between training and evaluation data, the addition of individual-level BCR properties resulted in extra accuracy gain compared with disease-specific properties (Figure 3E), thus demonstrating that BCR-SORT is applicable for chronological analysis of an individual’s B cell population with personalized optimization.

### Cell subset-aware lineage reconstruction

Beyond characterization of individual B cells, we demonstrated that BCR-SORT can also be applied to investigate maturation hierarchy within families of relevant B cells. As a proxy of maturation hierarchy, SHM hierarchy was inferred starting from SHM-free BCR sequences (germline) by simulating the accumulation of SHMs. To postulate the origin of such hierarchy, previous methods assume a virtual naïve B cell ancestor containing a germline sequence(39,40). However, the germline sequence cannot be specified around HCDR3 owing to unreliable alignment of IGHD germline gene, in other words, SHMs induced within HCDR3 cannot be fully elucidated. Consequently, previous methods are restricted in representing the actual B cell maturation hierarchy, and at times, result in an evolutionary scenario that contradicts the biology of B cell differentiation (e.g. ASC preceding naïve B cell).

This uncertainty in virtual naïve B cell ancestor is mitigated by BCR-SORT as it provides an actual naïve B cell as an alternative ancestor to replace the virtual one. Herein, using the alternative naïve B cell ancestor as a new root of the tree, we demonstrated that BCR-SORT contributes to rearranging the original phylogenetic tree, thereby suggesting alternative evolutionary scenarios better aligned with the biology of B cell differentiation (Figures 4A, S3). To this end, we designed a process to rearrange the phylogenetic tree inferred by previous methods (“reroot”) and evaluated the outcome of the process (Materials and methods). We compared the sequential order of BCR evolutions between phylogenetic trees reconstructed by IgPhyML and BCR-SORT via rank correlation (Figure 4B). Rerooting using BCR-SORT substantially altered the sequential order of BCR evolution, with the rerooting results well-aligned with those obtained using B cell subset information verified by FACS or scRNA-seq. In detail, rerooting using BCR-SORT relocated naïve B cells to the front of the tree, and ASCs to the rear (Figure 4C). Likewise, IgM BCRs were relocated to the front of the tree, whereas class-switched isotypes, such as IgG or IgA, were relocated to the rear (Figure 4D).

**Figure 4.**
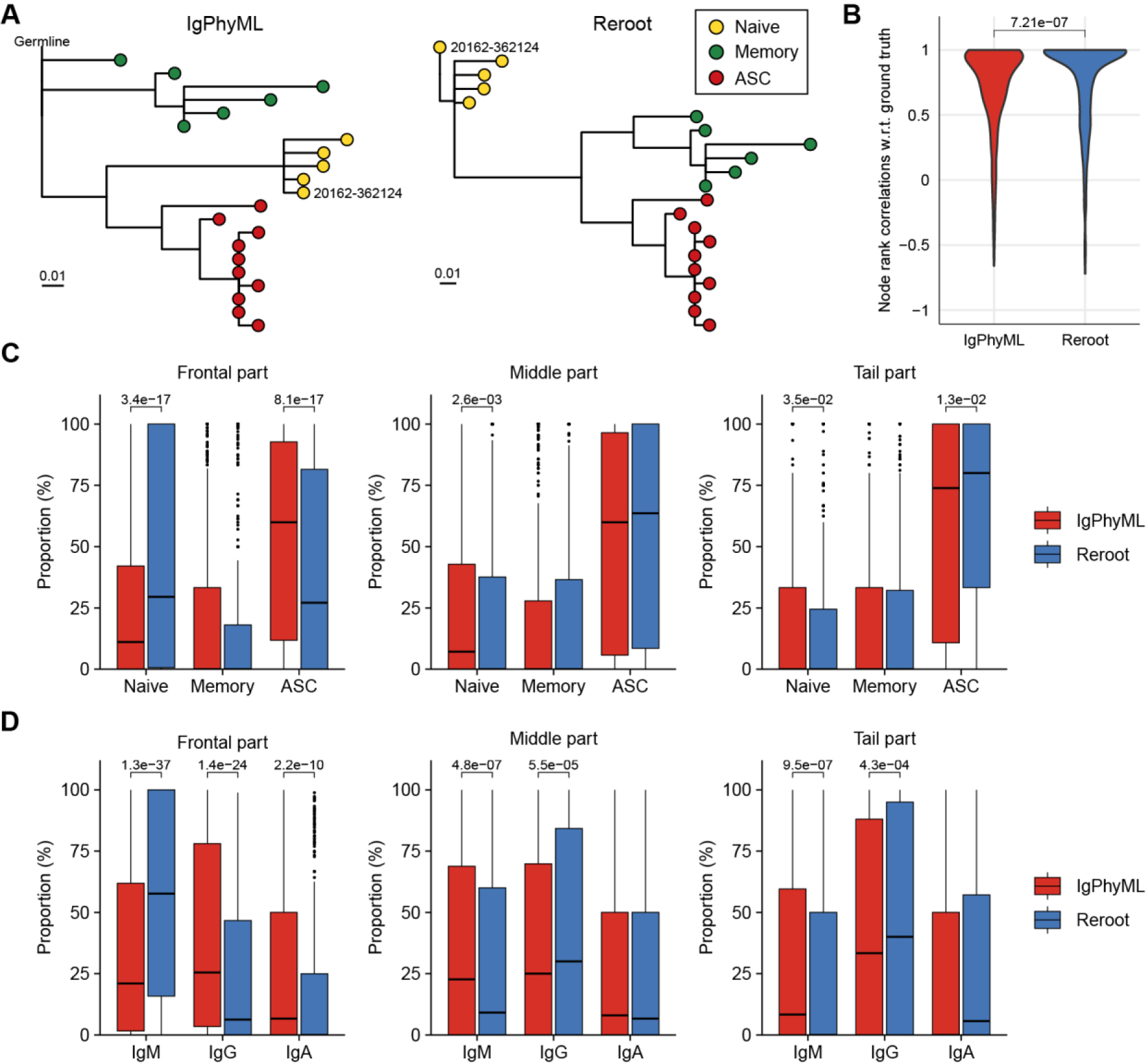
Comparisons of BCR phylogenetic trees inferred using cell subset-aware lineage reconstruction and conventional method. (**A**) An example of phylogenetic tree reconstructed using IgPhyML (left) and BCR-SORT (right). Naïve B cell node selected as a new root of the tree is labeled. (**B**) Distribution of rank correlations representing the sequential alignment of BCR evolutions between two phylogenetic trees. Within the same lineage, sequential order of BCR evolutions is identified along the phylogenetic tree reconstructed by IgPhyML and the tree rerooted by BCR-SORT, and compared with the tree rerooted based on the ground truth naïve B cells. (**C**, **D**) Effect of rerooting using BCR-SORT on rearranging BCRs within the phylogenetic tree. The distribution of B cell subset (**C**) and isotype (**D**) within the lineage is measured before and after rerooting using BCR-SORT. Each lineage is divided into three parts based on the evolutionary time scale, and positional variation of BCRs within the lineage is measured. In total, 442 lineages are analyzed in (**B**-**D**). Statistics are calculated using Mann-Whitney U test with p-value adjustment using Bonferroni correction in (**C**) and (**D**).

Owing to uncertainty of germline sequence, previous methods had difficulty in clarifying the BCR sequence development along evolution trajectory (affinity maturation process). On the other hand, lineage rerooting using BCR-SORT yielded clear mutation history since the root of the mutation was specified by the designated naïve B cell (Figure S4).

### Unveiling treatment-resistant B cell subpopulations in autoimmune disorders using BCR-SORT

Before applying BCR-SORT on publicly accessible, unlabeled BCR repertoire datasets, we first applied the model on in-house datasets. While these datasets were also unlabeled, they were anticipated to exhibit discernable patterns in the distribution of B cell subpopulations, thereby serving as a robust foundation for our preliminary assessments. Rituximab (RTX), a monoclonal antibody that targets CD20 molecules on B cells, is a treatment option for various autoimmune diseases mediated by autoantibody-producing B cells. RTX treatment induces profound alterations in B cell subpopulations by depleting naïve B cells and memory B cells, whereas ASCs are resistant to depletion owing to low CD20 expression on cell surface(55). Besides, these RTX-resistant ASCs have been substantiated to serve as a basis of post-RTX relapse(20). Thus, we utilized BCR-SORT to identify RTX-driven alterations and to investigate RTX-resistant ASCs with potential relevance to disease relapse.

We recruited one MG patient and one PV patient to investigate their BCR repertoires. Using BCR-SORT, we identified alterations in B cell subpopulations and observed that ASCs exhibited resistance to RTX treatment in both MG and PV patients (Figure 5A). As post-RTX relapse was identified in PV patient, autoantibody-producing B cells which resist the RTX treatment and survive until the relapse might be the potential source of the relapse. Therefore, we identified persistent clones, groups of similar BCR sequences that were present in all three peripheral blood sampling points (pre-RTX treatment, after it, and after disease relapse), and analyzed using BCR-SORT. Consistent with previous studies(20), ASCs defined by BCR-SORT appeared to be more persistent than memory B cells (Figure 5B). Moreover, phylogenetic analysis revealed that these persistent BCRs continuously acquired SHMs from pre-RTX stage to relapse, implying the continuous exposure to autoantigens (Figure 5C).

**Figure 5.**
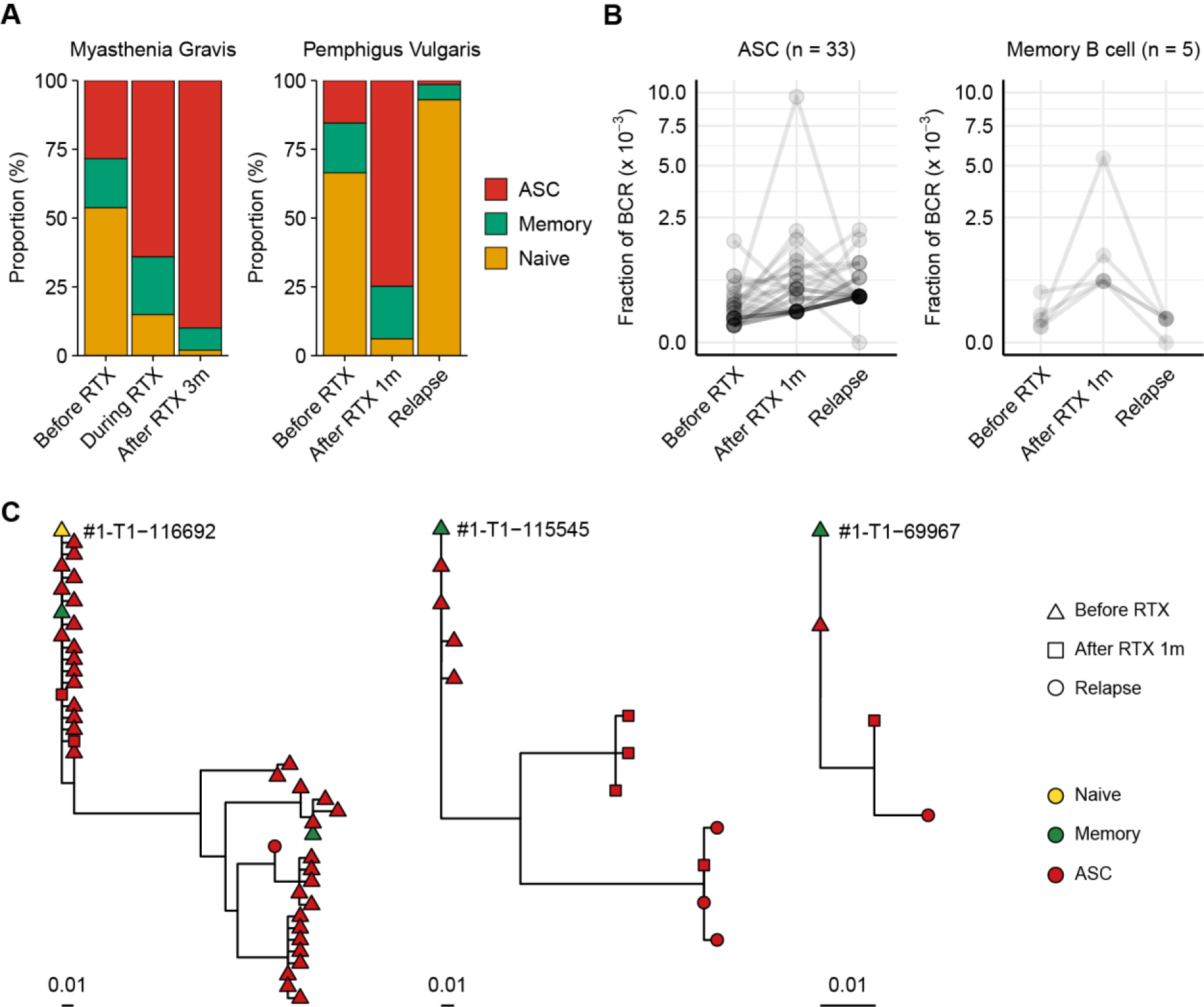
Chronological tracing of treatment-resistant B cells from autoimmune disease patients using BCR-SORT. (**A**) Proportion of B cell subsets throughout the course of treatment and relapse predicted using BCR-SORT. Data from MG patient (left) and PV patient (right) were shown. (**B**) BCR fractions in each persistent clone. Persistent clones were divided into those dominated by ASCs (left) and memory B cells (right), and their BCR fractions were measured within each subset. The number of persistent clones analyzed was indicated on top. (**C**) Representative phylogenetic tree of persistent clones accompanying multiple B cell subsets during the course of treatment and relapse.

### Elucidating COVID-19 vaccine-induced maturation of BCRs using BCR-SORT

Finally, we applied BCR-SORT to an unlabeled BCR repertoire dataset to interpret the role of B cells by recovering the B cell subset information. Using datasets constructed by Park et al.(48), BCR repertoires obtained from 41 COVID-19 vaccine recipients during three doses of vaccination were analyzed using BCR-SORT after fine-tuning the model with labeled COVID-19 data (Figure S5A). Prior to employing BCR-SORT, the intricate B cell responses against SARS-CoV-2 were not clearly understood (Figure S5B). However, following its use, these responses showed clear patterns distinct to each B cell subpopulation in accordance with the vaccination schedule (Figure 6A), representing the initial antigenic encounter (Dose 1), boosting (Dose 2), and the persistence of immunological memory over time (Dose 2+6wk and Dose 2+30wk), followed by rapid recall response (Dose 3).

**Figure 6.**
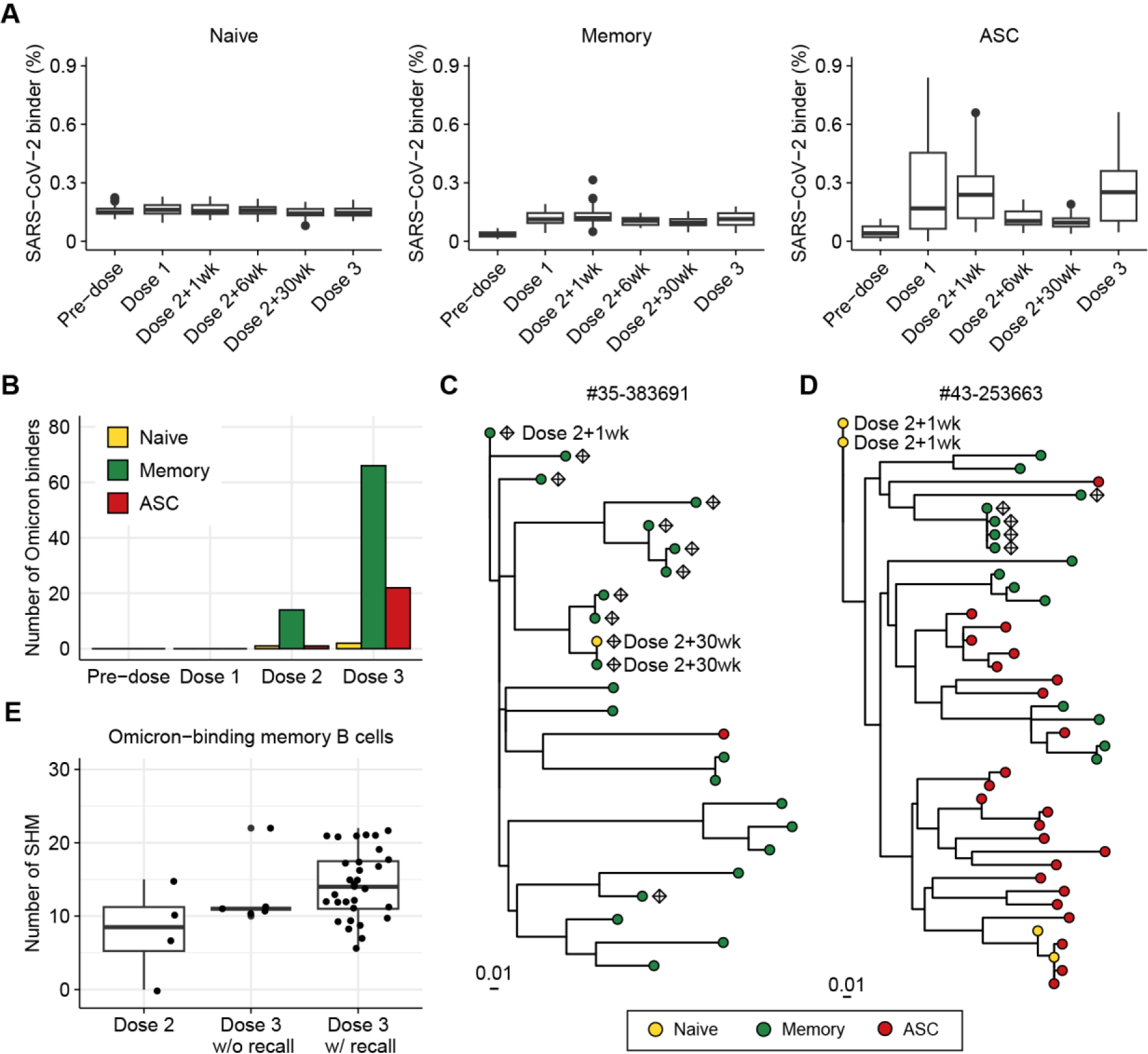
Investigation of BCR responses over triple COVID-19 vaccinations using BCR-SORT. (**A**) Proportion of SARS-CoV-2-binding BCRs with regard to B cell subsets predicted by BCR-SORT. For each vaccine recipient, BCR repertoires are identified at six different time points: pre-vaccination, 1 week after the first dose, 1, 6, 30 weeks after the 2^nd^ dose, and 1-4 weeks after the 3^rd^ dose. Outliers lying outside 2 standard deviations from the mean are discarded for better visualization. (**B**) B cell subset of Omicron-binding BCRs identified along multiple vaccinations. Omicron-binding BCRs are identified after the 2^nd^ dose and expand after the 3^rd^ dose, with them being predominantly predicted as memory B cells. (**C**, **D**) Phylogenetic tree of lineages containing Omicron-binding memory B cells obtained from two different vaccine recipients, with recall response from memory B cells originating from the 2^nd^ dose of vaccine (**C**), and without recall response (**D**). Sampling points are labeled for BCRs only when they are obtained except from the Dose 3, and Omicron-binding BCRs are labeled as diamond. (**E**) Omicron reactivity of BCRs emerging after the 2^nd^ dose and the 3^rd^ dose of vaccines. Verified EC50 values against Omicron viral proteins were shown for BCRs within lineage in (**C**) (left) and (**D**) (right).

Previous works have reported that triple vaccinations with the wild-type virus protein (original Wuhan strain) induces a potent antibody response to variants of concern including Omicron(56–58). Specifically, Omicron-binding memory B cells are known to emerge after the 2^nd^ dose of vaccine(59–61) and increase in number by the 3^rd^ dose of vaccine upon reactivation(56–58). However, the underlying landscape of SHMs and inter-individual heterogeneity of the recall response has remained unknown.

Consistent with the previously published findings, immunity against Omicron was found to be initiated by memory B cells defined by BCR-SORT after the 2^nd^ dose of vaccine, followed by their expansion after the 3^rd^ dose, with a large margin compared to ASCs (Figures 6B, S6). Of note, we identified three lineages from two vaccine recipients that contained Omicron-binding memory B cells, which exhibited significant expansion in number and evolutionary relationships between B cells emerging after the 2^nd^ and 3^rd^ doses. Using BCR-SORT, we identified inter-individual heterogeneity of memory B cell emergence during the course of vaccinations. Through phylogenetic analysis, we derived direct evidence that memory B cells emerging after the 2^nd^ dose of vaccines became the source of Omicron-specific recall response after the 3^rd^ dose via accumulation of SHMs (Figure 6C). However, no evidence of antigen experience was observed until the 3^rd^ dose of vaccine in a lineage obtained from another vaccine recipient (Figure 6D), although both lineages comprised similar HCDR3 sequences sharing an identical IGHV gene (Figure S7). Interestingly, the emergence of Omicron-specific memory B cells we captured was in accordance with the presence of antibodies exhibiting high levels of SHM, further enhancing our observations (Figures 6E, S8). Altogether, our observation suggested that the evolution of memory B cells could reveal individual differences in vaccine efficacy against SARS-CoV-2 variants.

## Discussion

In this study, we proposed BCR-SORT, a deep learning model designed to predict B cell subsets based on the given BCR sequence. Contrary to conventional B cell subset identification methods such as FACS or scRNA-seq, BCR-SORT enabled the coupling of antigen receptors with B cell subsets using solely the sequencing modality.

Leveraging HCDR3 sequences as a reliable source of B cell subset-specific features, BCR-SORT outperforms the current state-of-the-art method. When full-length BCR sequences were used as inputs instead of HCDR3 sequences, we observed no improvement in performance, further indicating the importance of HCDR3 in cell subset prediction (Figure S9). Extensive analysis on the model behavior via IG revealed that BCR-SORT utilized B cell activation and maturation signatures encoded in the HCDR3 sequence during B cell subset prediction. BCR-SORT was validated to be generally applicable to BCR datasets from various types of disease, with disease or individual-specific fine-tuning further enhancing the performance. In addition, BCR-SORT, in conjunction with conventional phylogenetic analysis method, enabled cell subset-aware rearrangement of BCR lineages, which yielded more interpretable results following the biology of B cell differentiation. Finally, BCR-SORT was applied to unlabeled datasets obtained from autoimmune disease patients and COVID-19 vaccine recipients. Notably, BCR-SORT not only reproduced results consistent with previous knowledge, but also elucidated the varying effects of vaccines on constructing immunological memory among different vaccine recipients.

Current limitation of BCR-SORT is the absence of a method to analyze individual BCRs while incorporating the context of the BCR repertoire, i.e., similar BCRs originating from the same source are not considered during prediction. In fact, Miho et al.(62) discovered that the similarity relations between BCRs exhibit distinctive features depending on the B cell subset, as the biological mechanism to diversify BCRs differs between cell subsets. Therefore, incorporating this information into predictions is expected to further improve the performance of the model. Additionally, BCR-SORT is unable to distinguish two different B cell subsets encoded by the identical BCR. Unfortunately, indistinguishable cases may emerge when the maturation of the B cell subset precedes BCR sequence maturation by a significant margin. However, results from post-hoc analysis on our dataset implied that these could be extremely rare in practice. We found that only 0.32%, 1.01%, and 0.09% of BCRs were found to be indistinguishable between naïve-memory, memory-ASC, and naïve-ASC compartments within our dataset, respectively. Finally, benchmarking scRNA-seq has been constrained due to limited datasets available compared to FACS, indicating the need for more extensive validation.

In recent developments, the integration of large language models (LLMs) has uncovered unknown representations within biological sequences, paving the way to harness their potential uses(63,64). By embracing the development of LLMs, complicated representations encoded in the HCDR3 sequences are expected to be further deciphered.

Lately, various analytic solutions have been proposed to decipher B cell or T cell responses by combining functional landscape and antigen specificity(65–69). However, these solutions focus on developing a novel method to integrate antigen receptor sequence data and gene expression data, while delegating the data generation to the costly scRNA-seq analysis. Consequently, these solutions are inherently restrained to be widely applied in various contexts and to cover high diversity of immune cells.

Instead, BCR-SORT focuses on the generation of missing B cell subset information to combine functional landscape and antigen specificity. High-throughput sequencing has already been established as a standard method to investigate BCR repertoire. Therefore, BCR-SORT is broadly applicable to establish missing links between individual BCR and cell subset. We anticipate that BCR-SORT will contribute to the generalization of the simultaneous analysis of B cell subsets and corresponding antigen receptors to clarify the roles of B cells during various immunological challenges.

## Author Contributions

HL: Conceptualization, Methodology, Software, Investigation, Writing – original draft, Writing – review & editing, Visualization. KS: Conceptualization, Software, Investigation, Writing – original draft, Writing – review & editing, Visualization. YL: Conceptualization, Methodology, Software, Writing – review & editing. Soobin Lee: Software, Investigation, Writing – original draft, Writing – review & editing. Seungyoun Lee: Investigation, Writing – original draft, Writing – review & editing. EL: Investigation, Writing – original draft, Writing – review & editing. SWK: Resources, Writing – review & editing, Supervision. HYS: Resources, Writing – review & editing, Supervision. JHK: Resources, Writing – review & editing, Supervision. JC: Resources, Writing – original draft, Writing – review & editing, Supervision, Project administration, Funding acquisition. SK: Conceptualization, Resources, Writing – original draft, Writing – review & editing, Supervision, Project administration, Funding acquisition.

## Supplementary Materials

**Supplementary Figure 1.**
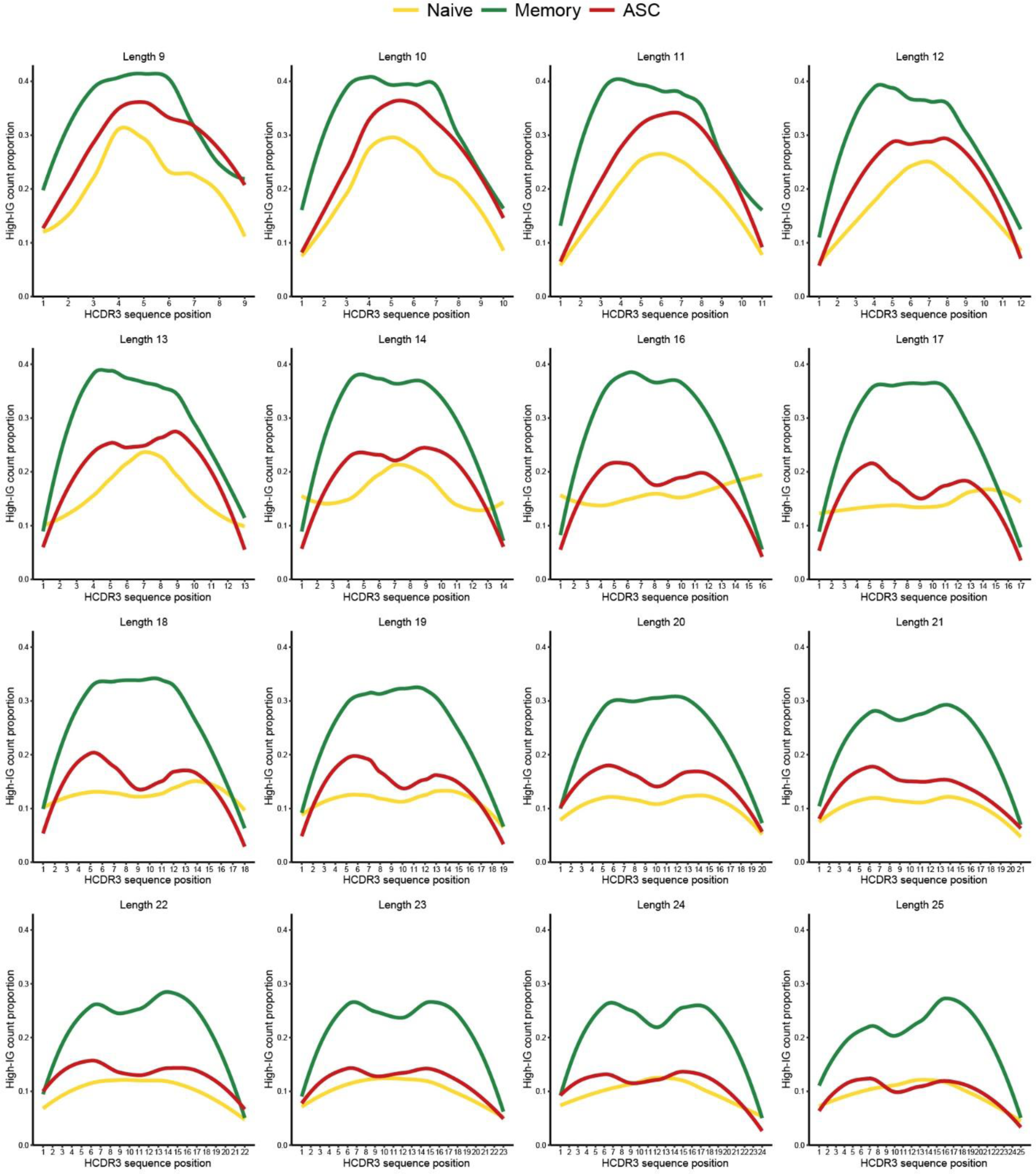
Proportion of high-IG HCDR3 residues with regard to position. All of the HCDR3 sequence lengths from 8 to 25 except 15 (which is shown in the main figure) are plotted.

**Supplementary Figure 2.**
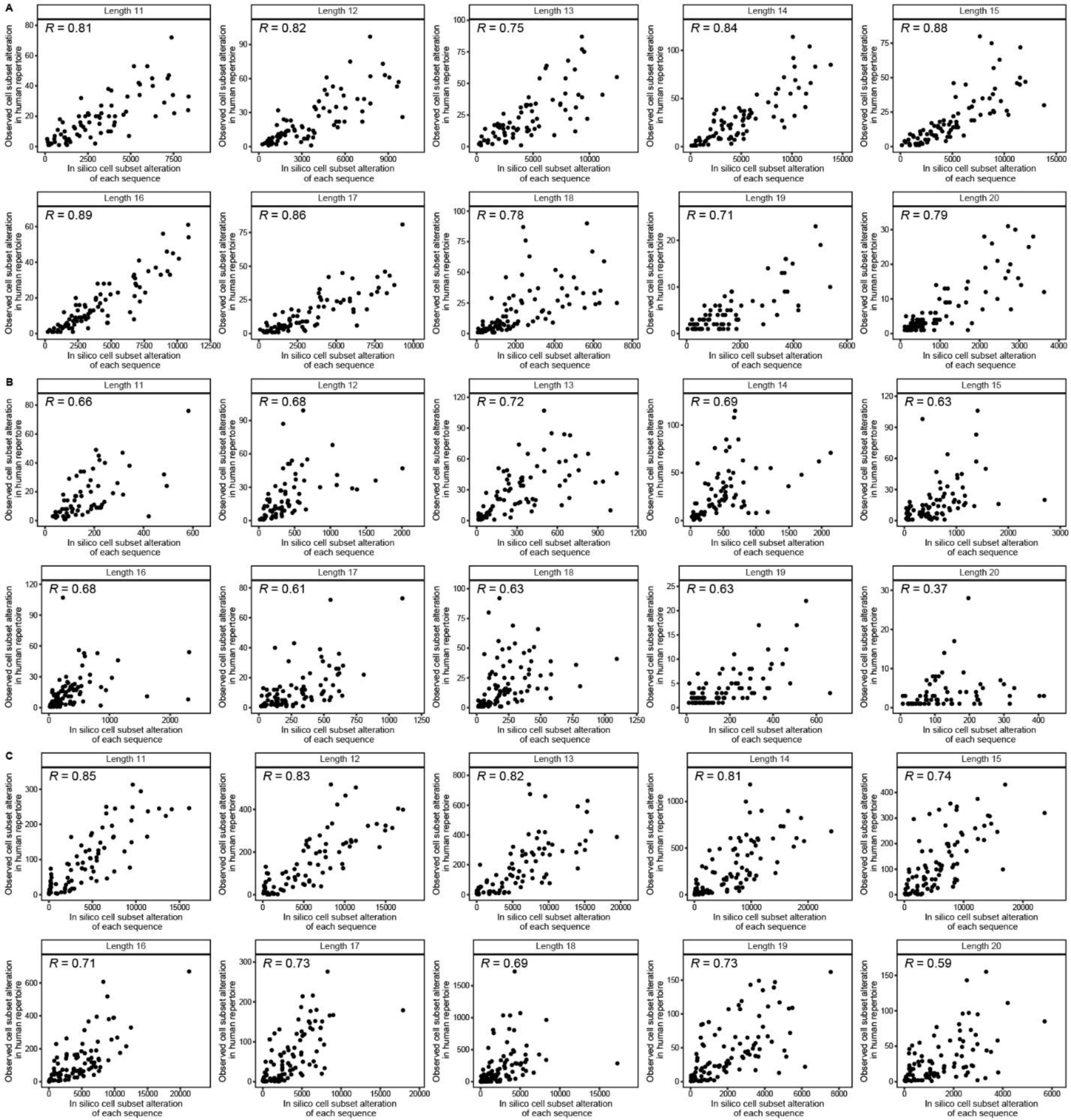
Correlation plots between the cell subset alteration counts measured in *in silico* saturation mutagenesis and in human repertoire data. Alteration of B cell subset between naïve-memory (**A**), memory-ASC (**B**), and naïve-ASC (**C**) are shown, respectively. Lengths between 11 and 20, which comprise the majority of the dataset, are plotted. Spearman’s rank correlation coefficient is shown in the figure.

**Supplementary Figure 3.**
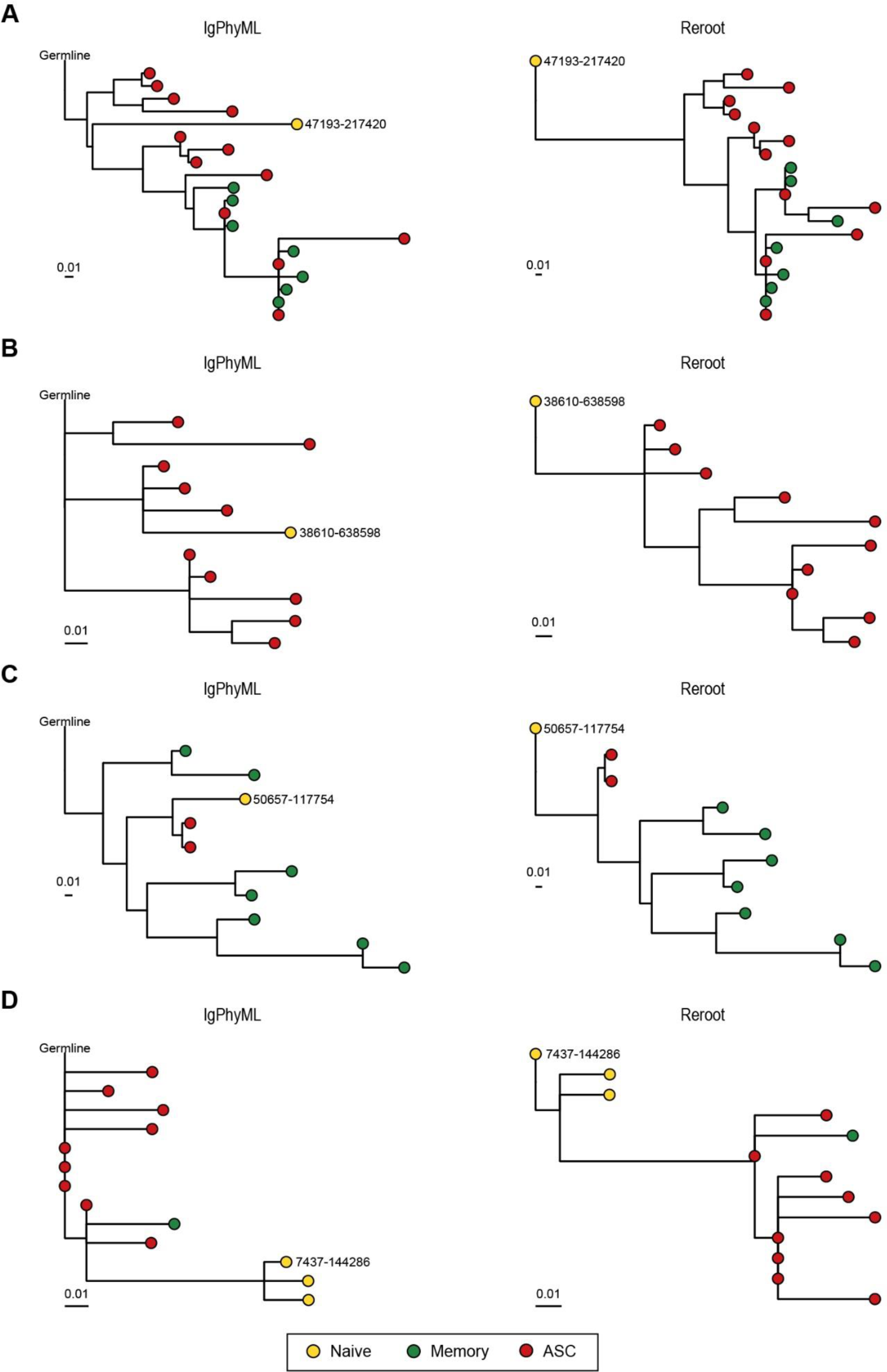
Visualization of BCR phylogenetic trees reconstructed using IgPhyML and BCR-SORT. Examples of phylogenetic trees originating from various diseases are illustrated to compare the two different methods. The type of disease analyzed corresponds to influenza vaccination (**A**), tetanus toxoid vaccination (**B**), healthy (**C**), and systemic lupus erythematosus (**D**). Naïve B cell node selected as a new root of the tree is labeled.

**Supplementary Figure 4.**
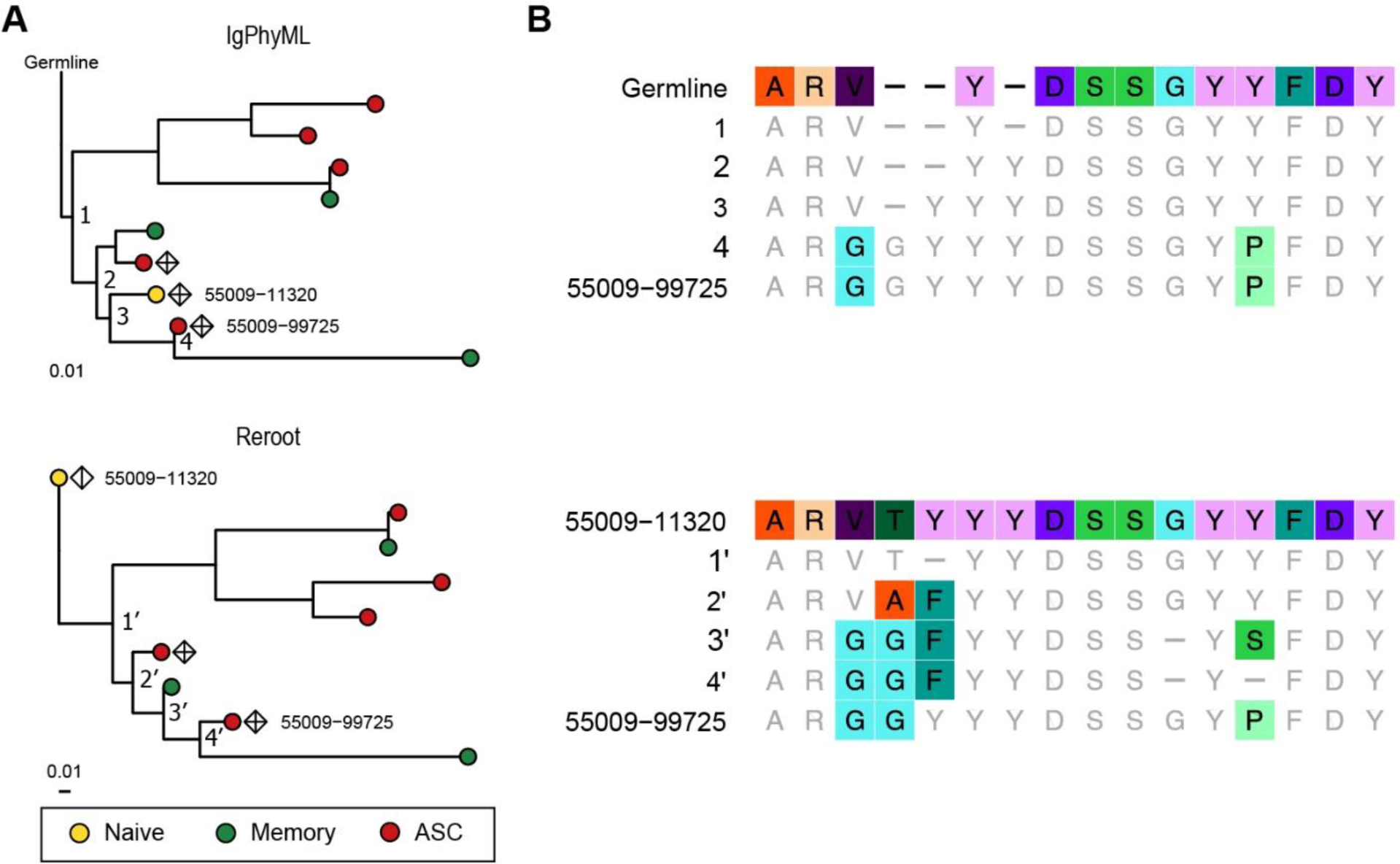
Rearranged mutation history toward antigen-binding BCRs. **(A)** An example of phylogenetic tree representing the maturation toward antigen-specific sequences. BCR lineage from COVID-19 patient is reconstructed by IgPhyML (top) and rerooted using BCR-SORT (bottom). Verified antigen-specific BCRs are labeled as diamond. **(B)** Alignment of BCR sequences along the maturation path in a. Sequence variations along with the maturation path defined by IgPhyML (top) and rerooted using BCR-SORT (down) are shown. Compared with IgPhyML, rerooting using BCR-SORT yields clearer HCDR3 sequence variations from the root sequence to the binder sequence. Amino acids are color-labeled when mutated from the prior sequence.

**Supplementary Figure 5.**
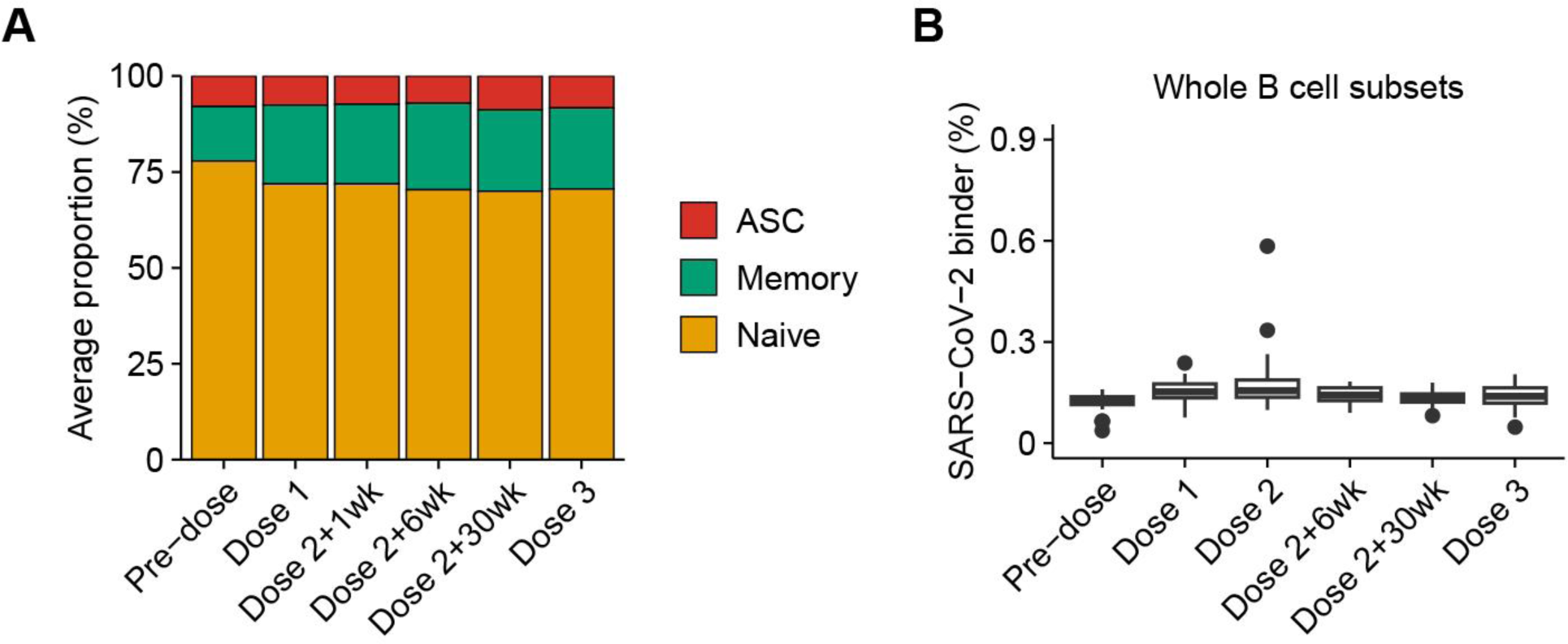
Chronological variation of B cell subpopulations during triple COVID-19 vaccinations. BCR repertoires are identified at six different time points: pre-vaccination, 1 week after the first dose, 1, 6, 30 weeks after the 2^nd^ dose, and 1-4 weeks after the third dose. (**A**) Proportion of B cell subsets constituting the BCR repertoires predicted using BCR-SORT. Average values over the entire COVID-19 vaccine recipient (n=41) are shown. (**B**) Proportion of SARS-CoV-2-binding BCRs without using B cell subset predicted by BCR-SORT. Outliers lying outside 2 standard deviations from the mean are discarded for better visualization.

**Supplementary Figure 6.**
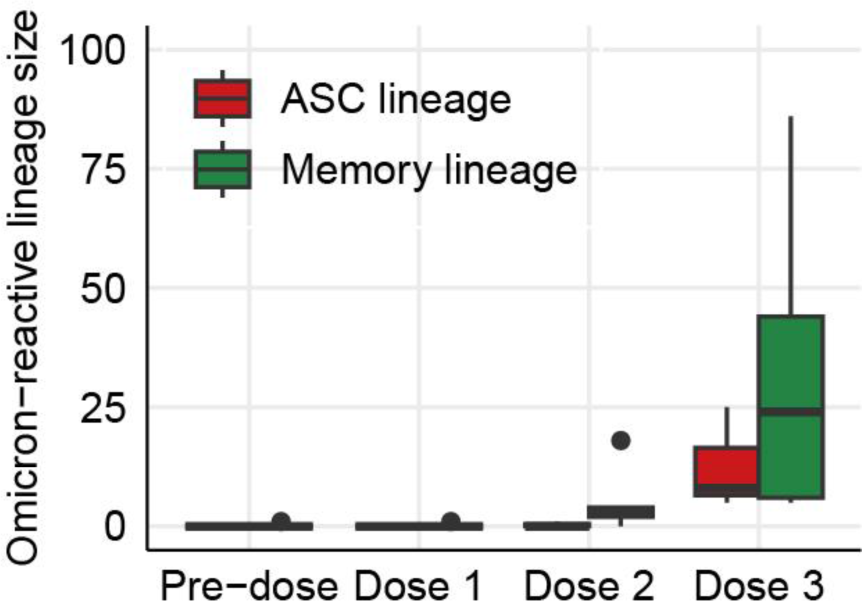
Expansion of BCR lineages containing Omicron binders during consecutive vaccinations. Size distribution of lineages containing Omicron-binding BCRs after the 3^rd^ dose. Lineage size is defined as the number of unique BCRs, and those containing five or more BCRs are considered (Memory lineage; n = 5, ASC lineage; n = 3).

**Supplementary Figure 7.**
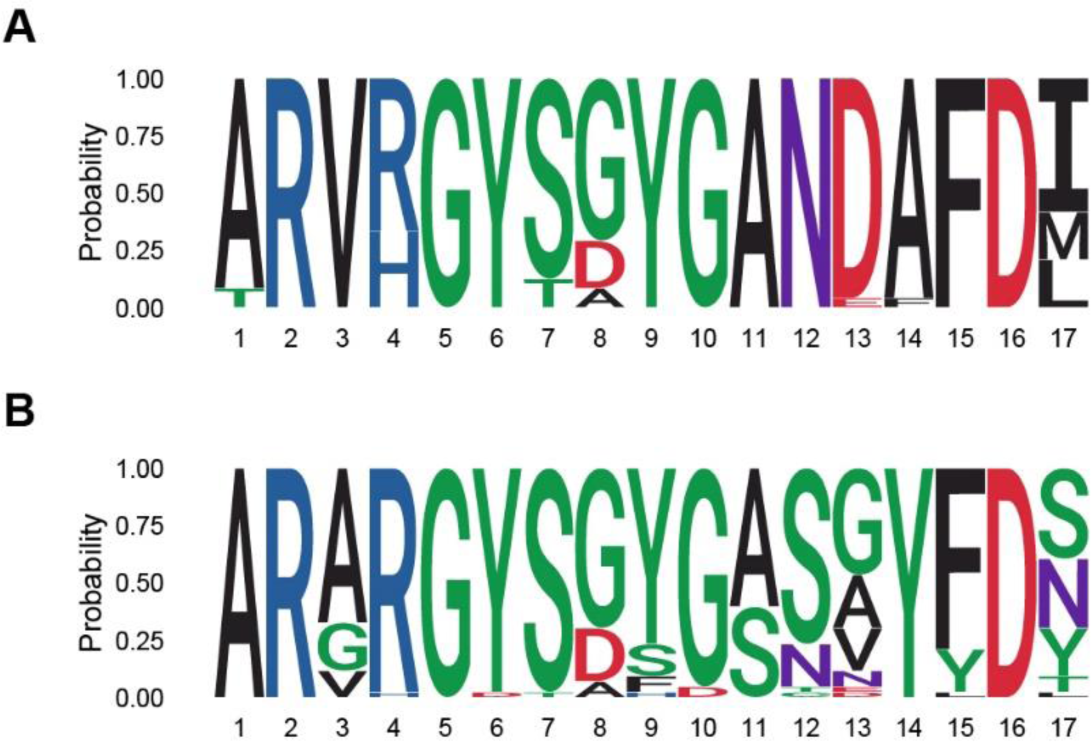
Logo plots visualizing HCDR3 sequences within Omicron-specific lineages. Lineage #35-383691 (**A**) and #43-253663 (**B**) are shown, respectively.

**Supplementary Figure 8.**
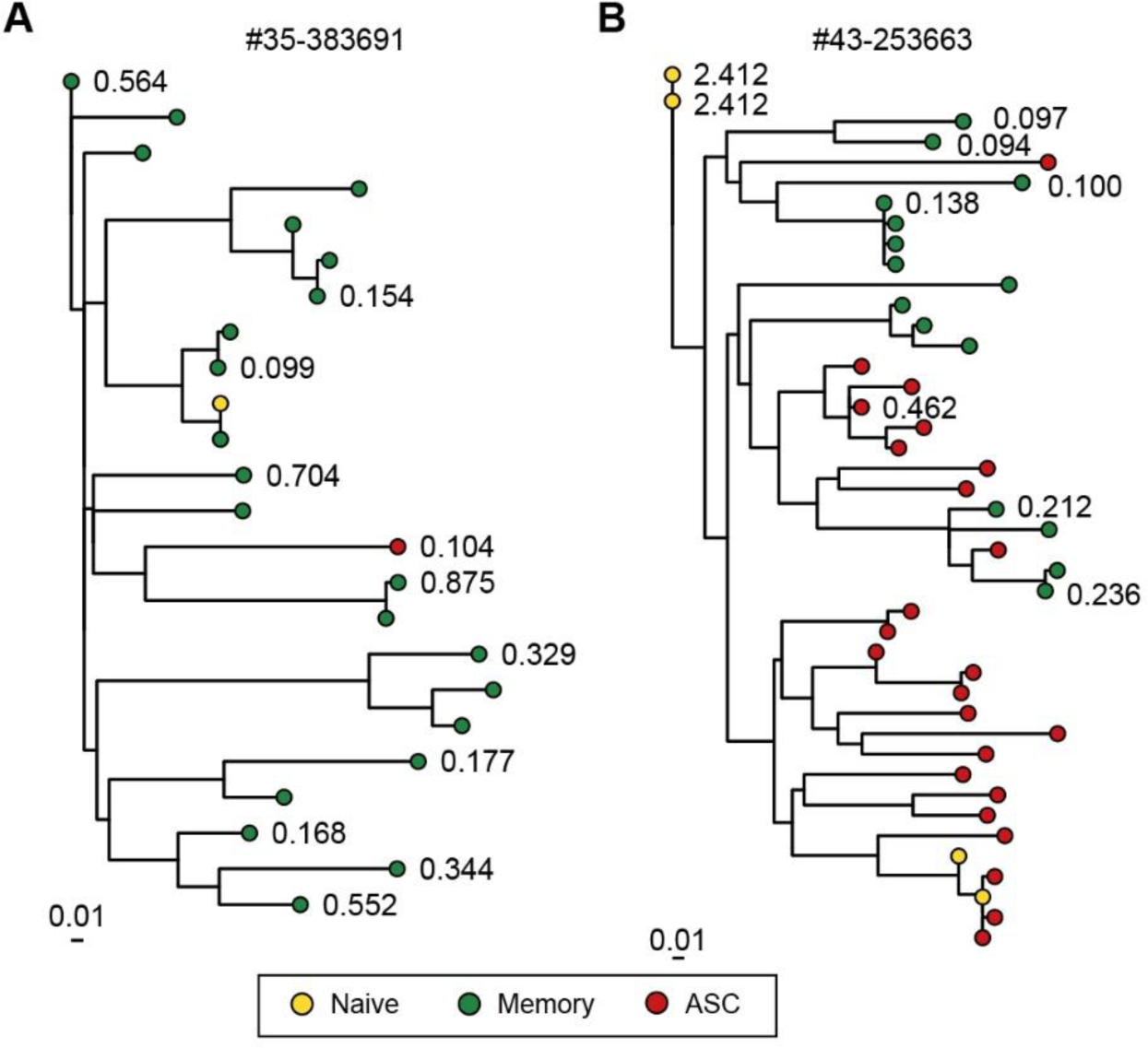
Omicron reactivity of BCRs comprising the Omicron-binding memory B cell lineages. BCRs with verified EC50 (nM) values against Omicron viral proteins were shown for lineage #35-383691 (**A**) and #43-253663 (**B**), respectively.

**Supplementary Figure 9.**
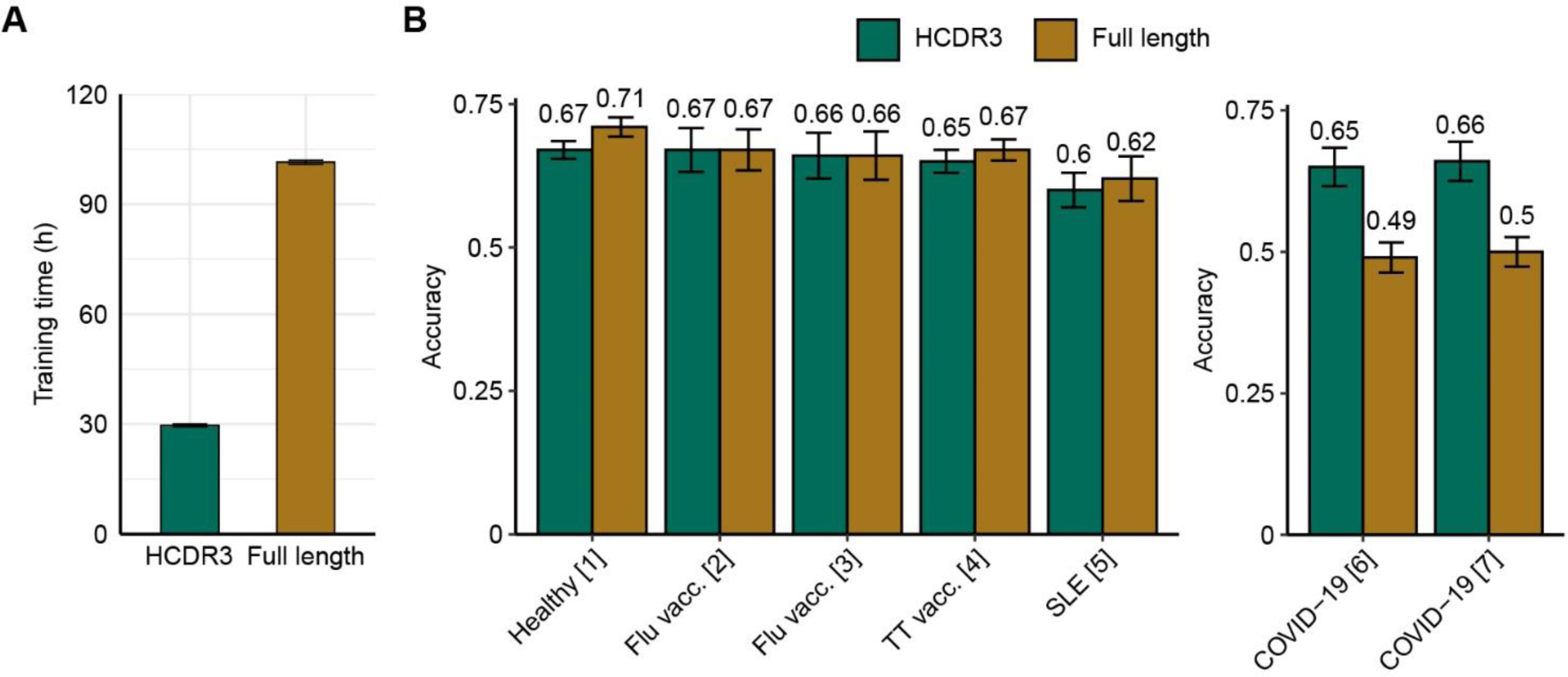
Comparison of HCDR3 and full-length model on benchmarking FACS and scRNA-seq. Instead of using the HCDR3 sequence, the full-length BCR is employed as the input for BCR-SORT, and their performances are compared. (**A)** Training time of BCR-SORT. (**B)** Performance of BCR-SORT on external validation datasets, verified by FACS (left) and scRNA-seq (right). Owing to the misalignment between full-length BCR sequences generated by FACS and scRNA-seq, the performance of BCR-SORT deteriorates when applied to scRNA-seq data, indicating that requiring full-length BCR reduces the model’s applicability. Accuracy is measured using various benchmark datasets and presented according to the source of the dataset. Performance is evaluated on datasets for whom full-length nucleotide BCR sequences are available, and error bars of datasets containing only a single sample are calculated by bootstrapping 150 BCR sequences 1,000 times. Although the full-length BCR sequence offers more information as input, the increase in performance is marginal considering the substantial computational cost incurred.

**Supplementary Table 1.**
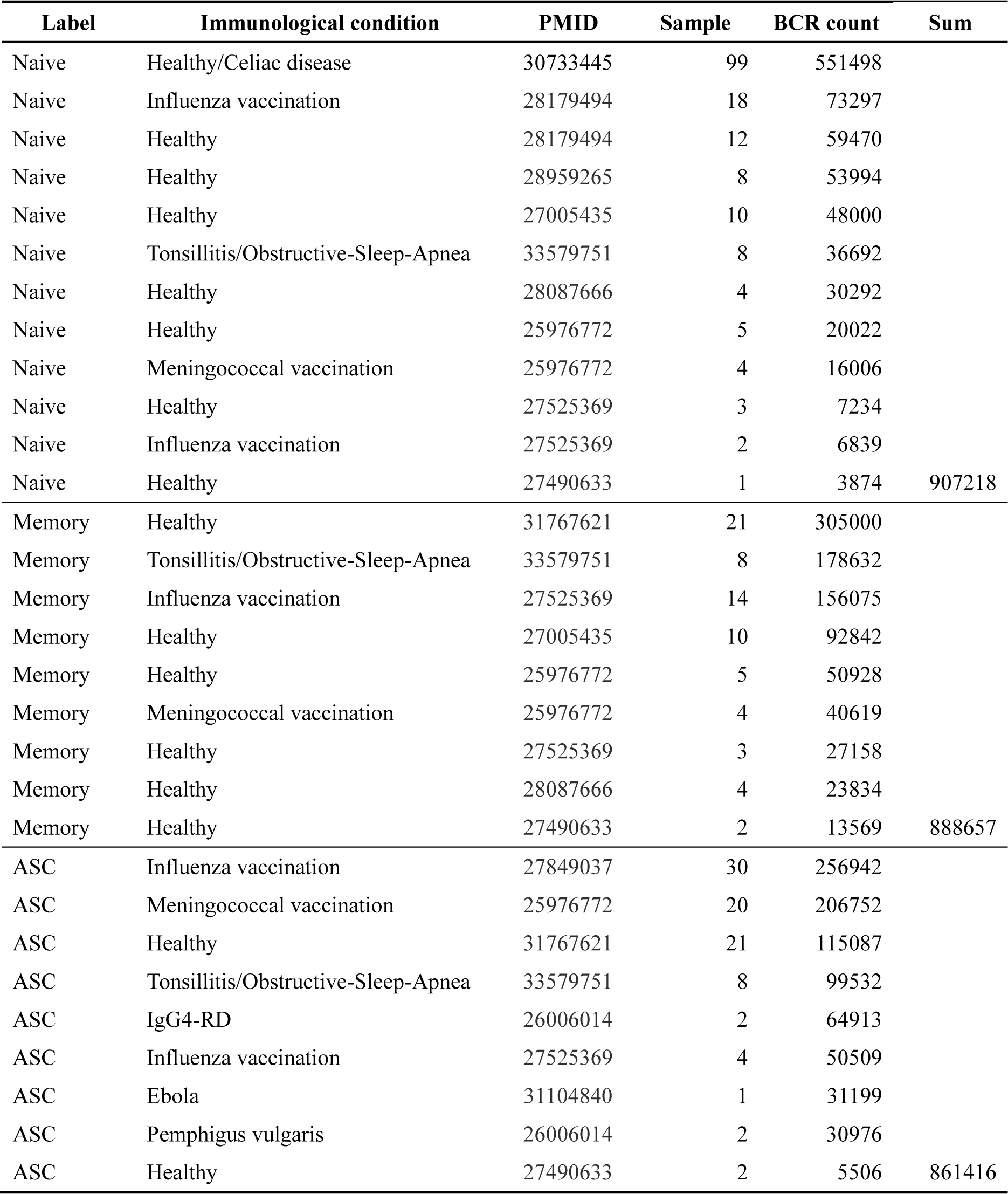
Statistics of datasets used for training.

| Label | Immunological condition | PMID | Sample | BCR count | Sum |
| --- | --- | --- | --- | --- | --- |
| Naive | Healthy/Celiac disease | 30733445 | 99 | 551498 |  |
| Naive | Influenza vaccination | 28179494 | 18 | 73297 |  |
| Naive | Healthy | 28179494 | 12 | 59470 |  |
| Naive | Healthy | 28959265 | 8 | 53994 |  |
| Naive | Healthy | 27005435 | 10 | 48000 |  |
| Naive | Tonsillitis/Obstructive-Sleep-Apnea | 33579751 | 8 | 36692 |  |
| Naive | Healthy | 28087666 | 4 | 30292 |  |
| Naive | Healthy | 25976772 | 5 | 20022 |  |
| Naive | Meningococcal vaccination | 25976772 | 4 | 16006 |  |
| Naive | Healthy | 27525369 | 3 | 7234 |  |
| Naive | Influenza vaccination | 27525369 | 2 | 6839 |  |
| Naive | Healthy | 27490633 | 1 | 3874 | 907218 |
| Memory | Healthy | 31767621 | 21 | 305000 |  |
| Memory | Tonsillitis/Obstructive-Sleep-Apnea | 33579751 | 8 | 178632 |  |
| Memory | Influenza vaccination | 27525369 | 14 | 156075 |  |
| Memory | Healthy | 27005435 | 10 | 92842 |  |
| Memory | Healthy | 25976772 | 5 | 50928 |  |
| Memory | Meningococcal vaccination | 25976772 | 4 | 40619 |  |
| Memory | Healthy | 27525369 | 3 | 27158 |  |
| Memory | Healthy | 28087666 | 4 | 23834 |  |
| Memory | Healthy | 27490633 | 2 | 13569 | 888657 |
| ASC | Influenza vaccination | 27849037 | 30 | 256942 |  |
| ASC | Meningococcal vaccination | 25976772 | 20 | 206752 |  |
| ASC | Healthy | 31767621 | 21 | 115087 |  |
| ASC | Tonsillitis/Obstructive-Sleep-Apnea | 33579751 | 8 | 99532 |  |
| ASC | IgG4-RD | 26006014 | 2 | 64913 |  |
| ASC | Influenza vaccination | 27525369 | 4 | 50509 |  |
| ASC | Ebola | 31104840 | 1 | 31199 |  |
| ASC | Pemphigus vulgaris | 26006014 | 2 | 30976 |  |
| ASC | Healthy | 27490633 | 2 | 5506 | 861416 |

**Supplementary Table 2.** Ablation study on input attributes.

| Methods | Accuracy |
| --- | --- |
| BCR-SORT | 0.833±0.002 |
| (w/o) CDR3 | 0.640±0.003 |
| (w/o) Isotype | 0.792±0.003 |
| (w/o) IGHV, IGHJ gene | 0.831±0.001 |

**Supplementary Table 3.**
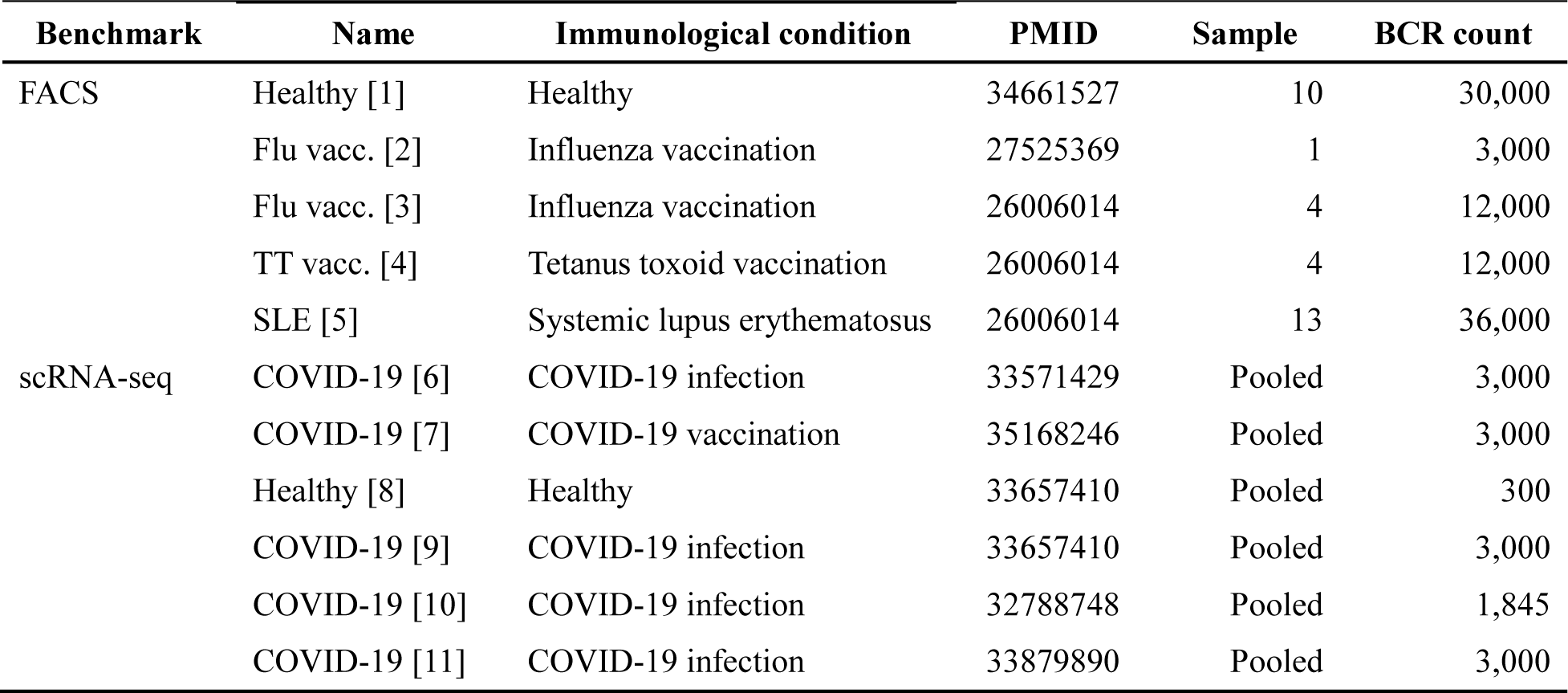
Statistics of datasets used for benchmarking FACS and scRNA-seq.

| **Benchmark** | **Name** | **Immunological condition** | **PMID** | **Sample** | **BCR count** |
| --- | --- | --- | --- | --- | --- |
| FACS | Healthy [1] | Healthy | 34661527 | 10 | 30,000 |
|  | Flu vacc. [2] | Influenza vaccination | 27525369 | 1 | 3,000 |
|  | Flu vacc. [3] | Influenza vaccination | 26006014 | 4 | 12,000 |
|  | TT vacc. [4] | Tetanus toxoid vaccination | 26006014 | 4 | 12,000 |
|  | SLE [5] | Systemic lupus erythematosus | 26006014 | 13 | 36,000 |
| scRNA-seq | COVID-19 [6] | COVID-19 infection | 33571429 | Pooled | 3,000 |
|  | COVID-19 [7] | COVID-19 vaccination | 35168246 | Pooled | 3,000 |
|  | Healthy [8] | Healthy | 33657410 | Pooled | 300 |
|  | COVID-19 [9] | COVID-19 infection | 33657410 | Pooled | 3,000 |
|  | COVID-19 [10] | COVID-19 infection | 32788748 | Pooled | 1,845 |
|  | COVID-19 [11] | COVID-19 infection | 33879890 | Pooled | 3,000 |

**Supplementary Table 4.** Isotype proportion of the dataset used for training.

| **Label** | **IgM (%)** | **IgD (%)** | **IgG (%)** | **IgA (%)** | **IgE (%)** |
| --- | --- | --- | --- | --- | --- |
| Naive | 77.236 | 19.424 | 2.453 | 0.883 | 0.005 |
| Memory | 54.527 | 1.703 | 31.047 | 12.719 | 0.004 |
| ASC | 9.196 | 0.000 | 60.739 | 30.044 | 0.021 |

## References

1. Hu X, Zhang J, Wang J, Fu J, Li T, Zheng X, Wang B, Gu S, Jiang P, Fan J, et al. Landscape of B cell immunity and related immune evasion in human cancers. Nat Genet (2019) 51:560–567. doi: 10.1038/s41588-018-0339-x

2. Wieland A, Patel MR, Cardenas MA, Eberhardt CS, Hudson WH, Obeng RC, Griffith CC, Wang X, Chen ZG, Kissick HT, et al. Defining HPV-specific B cell responses in patients with head and neck cancer. Nature (2021) 597:274–278. doi: 10.1038/s41586-020-2931-3

3. Biswas S, Mandal G, Payne KK, Anadon CM, Gatenbee CD, Chaurio RA, Costich TL, Moran C, Harro CM, Rigolizzo KE, et al. IgA transcytosis and antigen recognition govern ovarian cancer immunity. Nature (2021) 591:464–470. doi: 10.1038/s41586-020-03144-0

4. Ren X, Wen W, Fan X, Hou W, Su B, Cai P, Li J, Liu Y, Tang F, Zhang F, et al. COVID-19 immune features revealed by a large-scale single-cell transcriptome atlas. Cell (2021) 184:1895–1913.e19. doi: 10.1016/j.cell.2021.01.053

5. Sokal A, Chappert P, Barba-Spaeth G, Roeser A, Fourati S, Azzaoui I, Vandenberghe A, Fernandez I, Meola A, Bouvier-Alias M, et al. Maturation and persistence of the anti-SARS-CoV-2 memory B cell response. Cell (2021) 184:1201–1213.e14. doi: 10.1016/j.cell.2021.01.050

6. Stephenson E, Reynolds G, Botting RA, Calero-Nieto FJ, Morgan MD, Tuong ZK, Bach K, Sungnak W, Worlock KB, Yoshida M, et al. Single-cell multi-omics analysis of the immune response in COVID-19. Nat Med (2021) 27:904–916. doi: 10.1038/s41591-021-01329-2

7. Kim W, Zhou JQ, Horvath SC, Schmitz AJ, Sturtz AJ, Lei T, Liu Z, Kalaidina E, Thapa M, Alsoussi WB, et al. Germinal centre-driven maturation of B cell response to mRNA vaccination. Nature (2022) 604:141–145. doi: 10.1038/s41586-022-04527-1

8. Tonegawa S. Somatic generation of antibody diversity. Nature (1983) 302:575–581. doi: 10.1038/302575a0

9. Phad GE, Pinto D, Foglierini M, Akhmedov M, Rossi RL, Malvicini E, Cassotta A, Fregni CS, Bruno L, Sallusto F, et al. Clonal structure, stability and dynamics of human memory B cells and circulating plasmablasts. Nat Immunol 2022 (2022)1–10. doi: 10.1038/s41590-022-01230-1

10. Morgan D, Tergaonkar V. Unraveling B cell trajectories at single cell resolution. Trends Immunol (2022) 43:210–229. doi: 10.1016/j.it.2022.01.003

11. Kim S Il, Noh J, Kim S, Choi Y, Yoo DK, Lee Y, Lee H, Jung J, Kang CK, Song KH, et al. Stereotypic neutralizing VHantibodies against SARS-CoV-2 spike protein receptor binding domain in patients with COVID-19 and healthy individuals. Sci Transl Med (2021) 13:1–15. doi: 10.1126/scitranslmed.abd6990

12. Feldman J, Bals J, Altomare CG, Denis KS, Lam EC, Hauser BM, Ronsard L, Sangesland M, Moreno TB, Okonkwo V, et al. Naive human B cells engage the receptor binding domain of SARS-CoV-2, variants of concern, and related sarbecoviruses. Sci Immunol (2021) 6:1–13. doi: 10.1126/sciimmunol.abl5842

13. Hoffman W, Lakkis FG, Chalasani G. B cells, antibodies, and more. Clin J Am Soc Nephrol (2016) 11:137–154. doi: 10.2215/CJN.09430915

14. Cao Y, Su B, Guo X, Sun W, Deng Y, Bao L, Zhu Q, Zhang X, Zheng Y, Geng C, et al. Potent Neutralizing Antibodies against SARS-CoV-2 Identified by High-Throughput Single-Cell Sequencing of Convalescent Patients’ B Cells. Cell (2020) 182:73–84.e16. doi: 10.1016/j.cell.2020.05.025

15. Ju B, Zhang Q, Ge J, Wang R, Sun J, Ge X, Yu J, Shan S, Zhou B, Song S, et al. Human neutralizing antibodies elicited by SARS-CoV-2 infection. Nature (2020) 584:115–119. doi: 10.1038/s41586-020-2380-z

16. Rogers TF, Zhao F, Huang D, Beutler N, Burns A, He WT, Limbo O, Smith C, Song G, Woehl J, et al. Isolation of potent SARS-CoV-2 neutralizing antibodies and protection from disease in a small animal model. Science (80-) (2020) 369:956–963. doi: 10.1126/science.abc7520

17. Shi R, Shan C, Duan X, Chen Z, Liu P, Song J, Song T, Bi X, Han C, Wu L, et al. A human neutralizing antibody targets the receptor-binding site of SARS-CoV-2. Nature (2020) 584:120–124. doi: 10.1038/s41586-020-2381-y

18. Wu Y, Wang F, Shen C, Peng W, Li D, Zhao C, Li Z, Li S, Bi Y, Yang Y, et al. A noncompeting pair of human neutralizing antibodies block COVID-19 virus binding to its receptor ACE2. Science (80-) (2020) 368:1274–1278. doi: 10.1126/science.abc2241

19. Robbiani DF, Gaebler C, Muecksch F, Lorenzi JCC, Wang Z, Cho A, Agudelo M, Barnes CO, Gazumyan A, Finkin S, et al. Convergent antibody responses to SARS-CoV-2 in convalescent individuals. Nature (2020) 584:437–442. doi: 10.1038/s41586-020-2456-9

20. Jiang R, Fichtner ML, Hoehn KB, Pham MC, Stathopoulos P, Nowak RJ, Kleinstein SH, O’Connor KC. Single-cell repertoire tracing identifies rituximab-resistant B cells during myasthenia gravis relapses. JCI Insight (2020) 5: doi: 10.1172/JCI.INSIGHT.136471

21. Tipton CM, Fucile CF, Darce J, Chida A, Ichikawa T, Gregoretti I, Schieferl S, Hom J, Jenks S, Feldman RJ, et al. Diversity, cellular origin and autoreactivity of antibody-secreting cell population expansions in acute systemic lupus erythematosus. Nat Immunol (2015) 16:755–765. doi: 10.1038/ni.3175

22. Ellebedy AH, Jackson KJL, Kissick HT, Nakaya HI, Davis CW, Roskin KM, McElroy AK, Oshansky CM, Elbein R, Thomas S, et al. Defining antigen-specific plasmablast and memory B cell subsets in human blood after viral infection or vaccination. Nat Immunol (2016) 17:1226–1234. doi: 10.1038/ni.3533

23. Ghraichy M, Von Niederhausern V, Kovaltsuk A, Galson JD, Deane CM, Truck J. Different b cell subpopulations show distinct patterns in their igh repertoire metrics. Elife (2021) 10:1–15. doi: 10.7554/eLife.73111

24. Mikelov A, Alekseeva EI, Komech EA, Staroverov DB, Turchaninova MA, Shugay M, Chudakov DM, Bazykin GA, Zvyagin I V. Memory persistence and differentiation into antibody-secreting cells accompanied by positive selection in longitudinal BCR repertoires. Elife (2022) 11:1–29. doi: 10.7554/eLife.79254

25. Mitsunaga EM, Snyder MP. Deep characterization of the human antibody response to natural infection using longitudinal immune repertoire sequencing. Mol Cell Proteomics (2020) 19:278–293. doi: 10.1074/mcp.RA119.001633

26. Sutermaster BA, Darling EM. Considerations for high-yield, high-throughput cell enrichment: fluorescence versus magnetic sorting. Sci Rep (2019) 9:1–9. doi: 10.1038/s41598-018-36698-1

27. Sanz I, Wei C, Jenks SA, Cashman KS, Tipton C, Woodruff MC, Hom J, Lee FEH. Challenges and opportunities for consistent classification of human b cell and plasma cell populations. Front Immunol (2019) 10:1–17. doi: 10.3389/fimmu.2019.02458

28. Glanville J, Kuo TC, Von Büdingen HC, Guey L, Berka J, Sundar PD, Huerta G, Mehta GR, Oksenberg JR, Hauser SL, et al. Naive antibody gene-segment frequencies are heritable and unaltered by chronic lymphocyte ablation. Proc Natl Acad Sci U S A (2011) 108:20066–20071. doi: 10.1073/pnas.1107498108

29. Ghraichy M, Galson JD, Kovaltsuk A, von Niederhäusern V, Pachlopnik Schmid J, Recher M, Jauch AJ, Miho E, Kelly DF, Deane CM, et al. Maturation of the Human Immunoglobulin Heavy Chain Repertoire With Age. Front Immunol (2020) 11:1–13. doi: 10.3389/fimmu.2020.01734

30. De Bourcy CFA, Lopez Angel CJ, Vollmers C, Dekker CL, Davis MM, Quake SR. Phylogenetic analysis of the human antibody repertoire reveals quantitative signatures of immune senescence and aging. Proc Natl Acad Sci U S A (2017) 114:1105–1110. doi: 10.1073/pnas.1617959114

31. Kovaltsuk A, Leem J, Kelm S, Snowden J, Deane CM, Krawczyk K. Observed Antibody Space: A Resource for Data Mining Next-Generation Sequencing of Antibody Repertoires. J Immunol (2018) 201:2502–2509. doi: 10.4049/jimmunol.1800708

32. Ye J, McGinnis S, Madden TL. BLAST: Improvements for better sequence analysis. Nucleic Acids Res (2006) 34:W6–W9. doi: 10.1093/nar/gkl164

33. Ye J, Ma N, Madden TL, Ostell JM. IgBLAST: an immunoglobulin variable domain sequence analysis tool. Nucleic Acids Res (2013) 41:W34–W40. doi: 10.1093/nar/gkt382

34. Gupta NT, Heiden JA Vander, Uduman M, Gadala-Maria D, Yaari G, Kleinstein SH. Change-O: a toolkit for analyzing large-scale B cell immunoglobulin repertoire sequencing data. Bioinformatics (2015) 31:3356–3358. doi: 10.1093/bioinformatics/btv359

35. Kokhlikyan N, Miglani V, Martin M, Wang E, Alsallakh B, Reynolds J, Melnikov A, Kliushkina N, Araya C, Yan S, et al. Captum: A unified and generic model interpretability library for PyTorch. arXiv Prepr arXiv200907896 (2020) doi: 10.48550/arXiv.2009.07896

36. Liberis E, Velickovic P, Sormanni P, Vendruscolo M, Lio P. Parapred: Antibody paratope prediction using convolutional and recurrent neural networks. Bioinformatics (2018) 34:2944–2950. doi: 10.1093/bioinformatics/bty305

37. Wilson PC, Bouteiller O de, Liu Y-J, Potter K, Banchereau J, Capra JD, Pascual V. Somatic hypermutation introduces insertions and deletions into immunoglobulin V genes. J Exp Med (1998) 187:59–70.

38. Iman M, Arabnia HR, Rasheed K. A Review of Deep Transfer Learning and Recent Advancements. Technologies (2023) 11:40. doi: 10.3390/technologies11020040

39. Hoehn KB, Vander Heiden JA, Zhou JQ, Lunter G, Pybus OG, Kleinstein SH. Repertoire-wide phylogenetic models of B cell molecular evolution reveal evolutionary signatures of aging and vaccination. Proc Natl Acad Sci U S A (2019) 116:22664–22672. doi: 10.1073/pnas.1906020116

40. Hoehn KB, Lunter G, Pybus OG. A phylogenetic codon substitution model for antibody lineages. Genetics (2017) 206:417–427. doi: 10.1534/genetics.116.196303

41. Jeusset L, Abdollahi N, Verny T, Armand M, De Septenville AL, Davi F, Bernardes JS. ViCloD, an interactive web tool for visualizing B cell repertoires and analyzing intraclonal diversities: application to human B-cell tumors. NAR genomics Bioinforma (2023) 5:lqad064.

42. Hoehn KB, Pybus OG, Kleinstein SH. Phylogenetic analysis of migration, differentiation, and class switching in B cells. PLoS Comput Biol (2022) 18:e1009885. doi: 10.1371/journal.pcbi.1009885

43. Hoehn KB, Turner JS, Miller FI, Jiang R, Pybus OG, Ellebedy AH, Kleinstein SH. Title: Human b cell lineages associated with germinal centers following influenza vaccination are measurably evolving. Elife (2021) 10:e70873. doi: 10.7554/eLife.70873

44. Yu G. Using ggtree to Visualize Data on Tree-Like Structures. Curr Protoc Bioinforma (2020) 69:e96. doi: 10.1002/cpbi.96

45. Zhang J, Kobert K, Flouri T, Stamatakis A. PEAR: A fast and accurate Illumina Paired-End reAd mergeR. Bioinformatics (2014) 30:614–620. doi: 10.1093/bioinformatics/btt593

46. Sievers F, Higgins DG. Clustal omega, accurate alignment of very large numbers of sequences. Methods Mol Biol (2014) 1079:105–116. doi: 10.1007/978-1-62703-646-7_6

47. Sievers F, Wilm A, Dineen D, Gibson TJ, Karplus K, Li W, Lopez R, McWilliam H, Remmert M, Söding J, et al. Fast, scalable generation of high-quality protein multiple sequence alignments using Clustal Omega. Mol Syst Biol (2011) 7: doi: 10.1038/msb.2011.75

48. Park S, Choi J, Lee Y, Noh J, Kim N, Lee J, Cho G, Kim S, Yoo DK, Kang CK. An ancestral vaccine induces anti-Omicron antibodies by hypermutation. bioRxiv (2023)2003–2023. doi: 10.1101/2023.03.15.532728

49. Raybould MIJ, Kovaltsuk A, Marks C, Deane CM. CoV-AbDab: The coronavirus antibody database. Bioinformatics (2021) 37:734–735. doi: 10.1093/bioinformatics/btaa739

50. Morbach H, Eichhorn EM, Liese JG, Girschick HJ. Reference values for B cell subpopulations from infancy to adulthood. Clin Exp Immunol (2010) 162:271–279. doi: 10.1111/j.1365-2249.2010.04206.x

51. Sundararajan M, Taly A, Yan Q. Axiomatic attribution for deep networks. 34th Int Conf Mach Learn ICML 2017 (2017) 7:5109–5118. doi: 10.48550/arXiv.1703.01365

52. Finn JA, Leman JK, Willis JR, Cisneros A, Crowe JE, Meiler J. Improving loop modeling of the antibody complementarity-determining region 3 using knowledge-based restraints. PLoS One (2016) 11:1–15. doi: 10.1371/journal.pone.0154811

53. Alam O. In silico saturation mutagenesis of cancer genes. Nat Genet (2021) 53:1275. doi: 10.1038/s41588-021-00940-w

54. Zhang JY, Wang XM, Xing X, Xu Z, Zhang C, Song JW, Fan X, Xia P, Fu JL, Wang SY, et al. Single-cell landscape of immunological responses in patients with COVID-19. Nat Immunol (2020) 21:1107–1118. doi: 10.1038/s41590-020-0762-x

55. Palanichamy A, Jahn S, Nickles D, Derstine M, Abounasr A, Hauser SL, Baranzini SE, Leppert D, von Büdingen H-C. Rituximab Efficiently Depletes Increased CD20-Expressing T Cells in Multiple Sclerosis Patients. J Immunol (2014) 193:580–586. doi: 10.4049/jimmunol.1400118

56. Muecksch F, Wang Z, Cho A, Gaebler C, Ben Tanfous T, DaSilva J, Bednarski E, Ramos V, Zong S, Johnson B, et al. Increased memory B cell potency and breadth after a SARS-CoV-2 mRNA boost. Nature (2022) 607:128–134. doi: 10.1038/s41586-022-04778-y

57. Wang K, Jia Z, Bao L, Wang L, Cao L, Chi H, Hu Y, Li Q, Zhou Y, Jiang Y, et al. Memory B cell repertoire from triple vaccinees against diverse SARS-CoV-2 variants. Nature (2022) 603:919–925. doi: 10.1038/s41586-022-04466-x

58. Goel RR, Painter MM, Lundgreen KA, Apostolidis SA, Baxter AE, Giles JR, Mathew D, Pattekar A, Reynaldi A, Khoury DS, et al. Efficient recall of Omicron-reactive B cell memory after a third dose of SARS-CoV-2 mRNA vaccine. Cell (2022) 185:1875–1887.e8. doi: 10.1016/j.cell.2022.04.009

59. Cho A, Muecksch F, Schaefer-Babajew D, Wang Z, Finkin S, Gaebler C, Ramos V, Cipolla M, Mendoza P, Agudelo M, et al. Anti-SARS-CoV-2 receptor-binding domain antibody evolution after mRNA vaccination. Nature (2021) 600:517–522. doi: 10.1038/s41586-021-04060-7

60. Goel RR, Painter MM, Apostolidis SA, Mathew D, Meng W, Rosenfeld AM, Lundgreen KA, Reynaldi A, Khoury DS, Pattekar A, et al. mRNA vaccines induce durable immune memory to SARS-CoV-2 and variants of concern. Science (80-) (2021) 374:abm0829. doi: 10.1126/science.abm0829

61. Kotaki R, Adachi Y, Moriyama S, Onodera T, Fukushi S, Nagakura T, Tonouchi K, Terahara K, Sun L, Takano T, et al. SARS-CoV-2 Omicron-neutralizing memory B cells are elicited by two doses of BNT162b2 mRNA vaccine. Sci Immunol (2022) 7:eabn8590. doi: 10.1126/sciimmunol.abn8590

62. Miho E, Roškar R, Greiff V, Reddy ST. Large-scale network analysis reveals the sequence space architecture of antibody repertoires. Nat Commun (2019) 10:1321. doi: 10.1038/s41467-019-09278-8

63. Olsen TH, Moal IH, Deane CM. AbLang: An antibody language model for completing antibody sequences. Bioinforma Adv (2022)0–7. doi: 10.1093/bioadv/vbac046

64. Leem J, Mitchell LS, Farmery JHR, Barton J, Galson JD. Deciphering the language of antibodies using self-supervised learning. Patterns (2022)100513. doi: 10.1016/j.patter.2022.100513

65. Schattgen SA, Guion K, Crawford JC, Souquette A, Barrio AM, Stubbington MJT, Thomas PG, Bradley P. Integrating T cell receptor sequences and transcriptional profiles by clonotype neighbor graph analysis (CoNGA). Nat Biotechnol (2022) 40:54–63. doi: 10.1038/s41587-021-00989-2

66. Zhang Z, Xiong D, Wang X, Liu H, Wang T. Mapping the functional landscape of T cell receptor repertoires by single-T cell transcriptomics. Nat Methods (2021) 18:92–99. doi: 10.1038/s41592-020-01020-3

67. An Y, Drost F, Theis F, Schubert B, Lotfollahi M. Jointly learning T-cell receptor and transcriptomic information to decipher the immune response. bioRxiv (2021)2006–2021. doi: 10.1101/2021.06.24.449733

68. Sidhom JW, Larman HB, Pardoll DM, Baras AS. DeepTCR is a deep learning framework for revealing sequence concepts within T-cell repertoires. Nat Commun (2021) 12:1–12. doi: 10.1038/s41467-021-21879-w

69. Zhang Z, Chang WY, Wang K, Yang Y, Wang X, Yao C, Wu T, Wang L, Wang T. Interpreting the B-cell receptor repertoire with single-cell gene expression using Benisse. Nat Mach Intell (2022) 4:596–604. doi: 10.1038/s42256-022-00492-6

